# The T3SS structural and effector genes of *Chlamydia trachomatis* are expressed in distinct phenotypic cell forms

**DOI:** 10.1101/2024.04.25.591156

**Authors:** Nicole A. Grieshaber, Cody Appa, Megan Ward, Alorah Grossman, Sean McCormik, Brendan S. Grieshaber, Travis Chiarelli, Hong Yang, Anders Omsland, Scott S. Grieshaber

**Affiliations:** University of Idaho; Virginia Commonwealth University School of Medicine; Washington State University

## Abstract

Bacteria in the chlamydiales order are obligate intracellular parasites of eukaryotic cells. Within this order, the genus *Chlamydia* contains the causative agents of a number of clinically important infections of humans. Biovars of *C. trachomatis* are the causative agents of trachoma, the leading cause of preventable blindness worldwide, as well as sexually transmitted infections with the potential to cause pelvic inflammatory disease and infertility. Irrespective of the resulting disease, all chlamydial species share the same obligate intracellular life cycle and developmental cell forms. They are reliant on an infectious cycle consisting of at least three phenotypically distinct cell forms termed the reticulate body (RB), the intermediate body (IB) and the elementary body (EB). The EB is infectious but does not replicate. The RB replicates in the host cell but is non-infectious, while the IB is an intermediate form that transitions to the EB form. In this study, we ectopically expressed the transcriptional repressor Euo, the two nucleoid-associated proteins HctA and HctB, and the two component sensor kinase CtcB in the RB. Transcriptional analysis using RNA-seq, differential expression clustering and fluorescence *in situ* hybridization analysis show that the chlamydial developmental cycle is driven by three distinct regulons corresponding to the RB, IB or EB cell forms. Moreover, we show that the genes for the T3SS were cell type restricted, suggesting defined functional roles for the T3SS in specific cell forms.

**Importance:** *Chlamydia trachomatis*, a sexually transmitted bacterial infection, poses a significant global health threat, causing over 100 million infections annually and leading to complications like ectopic pregnancy and infertility. This study investigates the gene expression patterns of *Chlamydia trachomatis* during its unique life cycle within human cells. As an obligate intracellular parasite, *C. trachomatis* transitions through distinct developmental stages - one for infection and dissemination, another for replication, and a third for transitioning back to the infectious form. By analyzing gene expression profiles at each stage, we identified key genes involved in these processes. Interestingly, our research also reveals the presence of two separate T3SS (Type III Secretion System) translocons expressed in distinct stages, suggesting their crucial roles in specific functions during the infection cycle.

## Introduction

Many bacterial species undergo dramatic phenotypic changes to adapt to different environments or to generate cells with specific physiological functions. All the bacteria in the Chlamydiales order are obligate intracellular parasites of eukaryotic cells that undergo a developmental cycle with both non-replicating and actively replicating cell forms [1,2]. Chlamydial species are important pathogens of humans. *C. psittaci* causes zoonotic infections resulting in pneumonia, while *C. pneumoniae* is a human pathogen that causes respiratory disease. Different biovars of *C. trachomatis (Ctr)* are the causative agents of trachoma, the leading cause of preventable blindness worldwide, as well as sexually transmitted infections with the potential to cause pelvic inflammatory disease, ectopic pregnancy and infertility [3–5].

Success of a chlamydial infection depends on the completion of a complex intracellular developmental cycle, consisting of multiple cell forms; the elementary body (EB), the reticulate body (RB) and the intermediate body (IB) [1,2]. Although the timing of cell type conversion may differ, the broad strokes of this cycle are conserved in all the Chlamydiaceae [6,7]. Our current understanding of the developmental cycle as determined through promoter reporter strains, single inclusion kinetics, single cell gene expression and agent based modeling has led to a clearer picture of the cycle [8,9]. The EB, characterized by its condensed nucleoid and small size (∼0.2 nm diameter), initiates infection of the host through the use of a Type III Secretion System (T3SS) and pre-formed effectors [10,11]. These effectors promote pathogen phagocytosis and entry into the targeted cell. After entry, the EB form resides in an endocytic vesicle termed the inclusion that is modified through chlamydial gene expression [10,11]. The EB completes EB to RB differentiation and becomes replication competent at ∼10 hpi (*Ctr* serovar L2) [12,13]. The RB, which is phenotypically characterized as larger than the EB (∼1 nm diameter) and containing a dispersed nucleoid, then undergoes several rounds of amplifying replication before maturing to produce IB cells that then progress to the infectious EB, a process that takes place over ∼8-10 hours after IB formation [8,9]. The mature RBs continue to produce IB cells, acting akin to a stem cell population [8]. This developmental program results in a growth cycle that does not act like a typical bacterial growth culture (lag, log, stationary phase) but instead asynchronously progresses through the RB, IB and EB cell type transitions until cell lysis or inclusion extrusion [8,9].

The current understanding of the regulation of the developmental cycle comes primarily from population-level studies that frame the cycle in terms of time, treating the chlamydial population as a time dependent uniform culture. Population level gene expression data has been determined for chlamydial infections and has been described according to time after infection. These studies include RT-qPCR, microarray and RNA-seq studies and contribute to the canonical early (∼0-10 hpi, EB to RB differentiation), mid-cycle (∼10-18 hpi, RB replication) and late (∼18 hpi-cell lysis, EB formation) gene expression paradigm [14–18]. The reliance on population level data from this mixed cell population has confounded the understanding of gene expression as it pertains to the specific chlamydial cell forms.

Here we sought to define the transcript profiles of the cell forms that underpin the observed growth cycle by investigating the effects of ectopic expression of four transcriptional regulatory proteins in *Chlamydia trachomatis* (*Ctr*): Euo, HctA, HctB and CtcB. Euo (Early Upstream ORF) is among the earliest genes expressed post EB to RB differentiation during chlamydial infection [19–21]. Current evidence suggests that Euo is a DNA binding protein that acts to repress a handful of late cycle genes [19–21] and Euo ectopic expression leads to a block in the developmental cycle [22]. HctA is a small DNA binding protein with limited homology to the histone H1 histone family and is expressed transiently in the IB cell type ∼8-10 hrs before HctB [8,9,23]. HctA has been shown to bind DNA and to repress transcription broadly across chromosomes of both *Ctr* and, when ectopically expressed, *E. coli* [23–25]. HctB is a second small positively charged protein that has limited homology to the H1 histone family and is thought to contribute to condensation of the EB nucleoid [26]. Our data show that unlike HctA, HctB is expressed late in EB development, during the final stages of EB formation [8,9]. In addition to these DNA binding proteins, *Ctr* contains a single cytosolic two component regulatory system (TCS) consisting of CtcB/CtcC (histidine kinase/response regulator) [27]. The *Ctr* TCS is actively transcribed during RB to EB development, and the protein products are functional with respect to phosphotransfer [28]. Additionally, expression of the ATPase effector domain of the response regulator, CtcC, resulted in the up regulation of the sigma54 regulon which included many developmentally regulated genes [29]. We expect that ectopic expression of CtcB would phosphorylate CtcC and amplify the signaling and activation process of sigma54 gene expression.

We have shown that Euo, HctA and HctB promoter activities help define the RB, IB and EB cell forms [8,9]. Therefore, along with CtcB we determined the effects of ectopic expression of these regulatory factors on the transcriptome of *Ctr* using RNA-seq. Our data produced gene regulation profiles consistent with cell form-specific transcriptomes. This allowed us to assign/predict the expression of a large fraction of the chlamydial genome into RB, IB and EB-specific transcript categories. Within our cell form expression prediction groups were a number of T3SS genes. Using fluorescence *in situ* hybridization (FISH) in the context of cell type promoters, we showed that components of the T3SS were expressed in specific cell forms.

## Results

### Ectopic expression of Euo, HctA, CtcB, and HctB resulted in arrest of the developmental cycle

To determine the effects of the ectopic expression of Euo, HctA, HctB and CtcB on gene expression and the developmental cycle, we expressed these proteins as well as the GFP protein Clover (control) under the control of the T5 promoter and theophylline responsive riboswitch from the native chlamydial plasmid [30]. Cells were infected with the strains L2-E-euo-FLAG, L2-E-hctA-FLAG, L2-E-ctcB-FLAG, L2-E-clover-FLAG and L2-tet-J-E-hctB-FLAG and protein expression was induced at 15 hpi. We chose to induce expression at 15 hpi in order to evaluate the effects of these proteins on the RB to EB stage of the developmental cycle. At 15 hpi the vast majority of the chlamydial cells would be in the RB form [8].

We tested for the production of infectious progeny (EBs) using a reinfection assay at 48 hpi. Ectopic expression of all four transcriptional regulatory proteins resulted in a significant inhibition of EB production as compared to the Clover-FLAG controls (Fig. 1A). In addition, infected cells were imaged using transmission electron microscopy (TEM). For TEM cells were infected with the four strains plus the Clover control strains, induced for expression at 15 hpi and fixed and prepared for TEM analysis at 30 hpi (Fig. 1B). The induced and uninduced Clover-FLAG control samples were indistinguishable. The inclusions for both samples had similar ratios of RBs (large cell forms) and EBs (small electron dense forms). Cells ectopically expressing Euo had inclusions with very few visible EBs (small electron dense forms) and most cells appeared RB like (large less electron dense cell forms). The inclusions for the HctB and HctA expressing bacteria contained small populations of abnormal RB like forms as well as cells with dense structures resembling condensed nucleoids (electron dense regions inside cells). The inclusions of the CtcB expressing bacteria contained both RB and EB-like cells and an increase in intermediate forms, i.e. RB sized cells with condensed nucleoids (Fig. 1B). These data suggested that ectopic expression of all four of these transcriptional regulatory proteins resulted in an aborted developmental cycle as indicated by both the IFU measurements and the dysregulated cell forms seen by EM.

**Figure 1.**
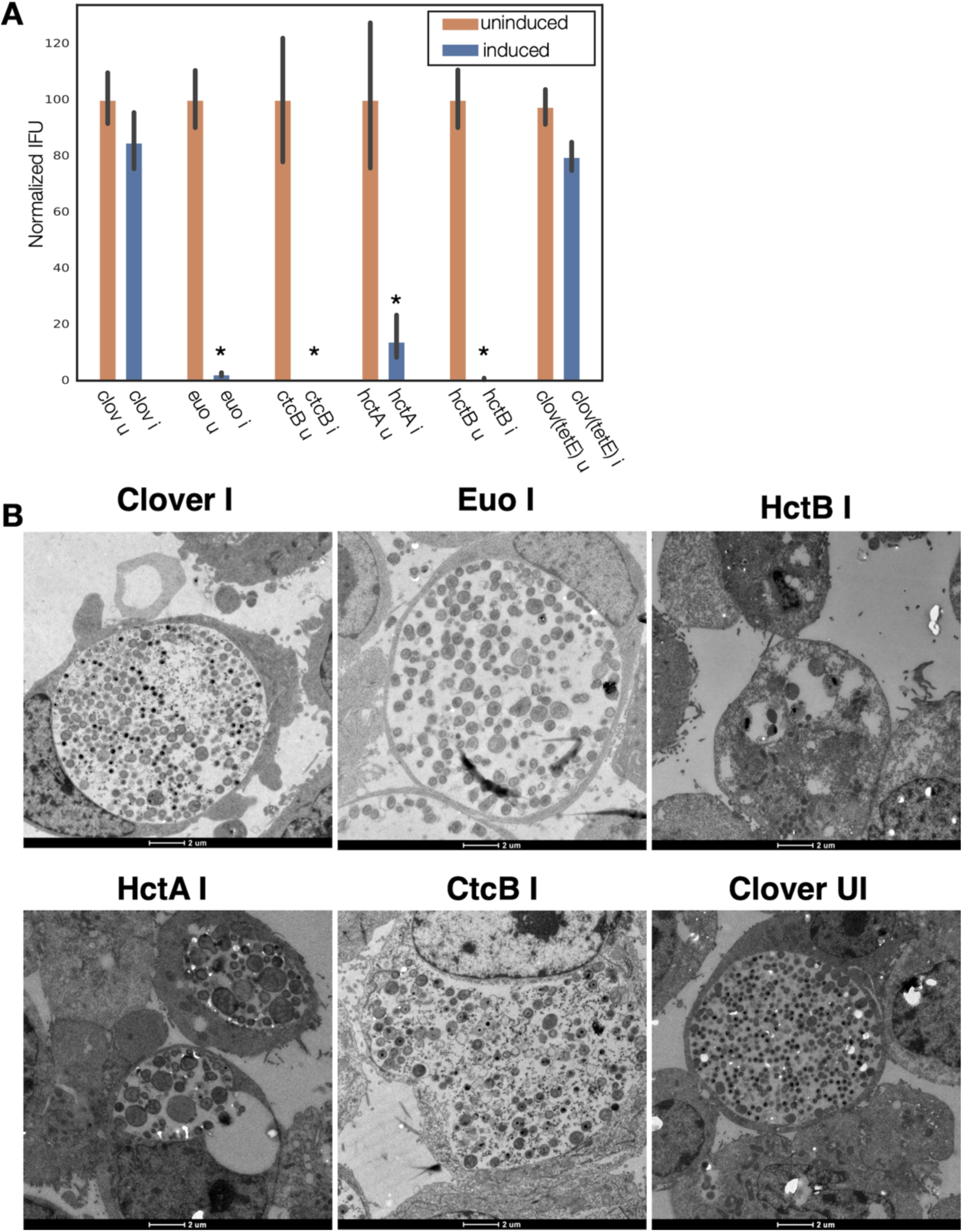
Ectopic expression of Euo, HctA, CtcB and HctB resulted in inhibition of the developmental cycle. (A) Cos-7 cells were infected with the four strains and a Clover control strain and induced for ectopic expression at 15 hpi. EBs were harvested at 48 hpi. IFU production was dramatically reduced by the ectopic expression of Euo, HctA, CtcB and HctB but not by the expression of the Clover protein. * = P < 0.01. (B) Transmission EM of Cos-7 cells infected with *Ctr* expressing Clover, Euo, HctB, HctA or CtcB. Ectopic expression was induced at 15 hpi and the cells were fixed and prepared for imaging at 30 hpi. The bacteria in the induced and uninduced Clover control chlamydial infections looked similar with inclusions of both samples containing large RB like cells as well as electron-dense EB-like cells. The *Ctr* in the Euo expressing inclusions were primarily RB like cells while very few cells were electron dense EB cells. The chlamydial cells in the HctB expressing inclusions were abnormal looking, some with apparent condensed nucleoids. The HctA expressing *Ctr* also appeared abnormal with condensed nucleoids. Many of the CtcB expressing *Ctr* cells were target-like RB sized cells with a condensed nucleoid.

### RNA-seq of the ectopically expressing chlamydial strains

To better understand the effects of the ectopic expression of each regulatory protein, we used RNA-seq to characterize the corresponding transcriptomes. We infected host cells with each strain (L2-E-clover-FLAG, L2-E-euo-FLAG, L2-E-hctA-FLAG, L2-E-ctcB-FLAG, L2-tet-J-E-hctB-FLAG and and L2-E-clover-FLAG), induced expression at 15 hpi and harvested RNA for library construction at 18 hpi and 24 hpi. We chose to investigate gene expression three hours after induction (18 hpi) to capture potential immediate effects on the developmental cycle. We also investigated gene expression at 9 hours after induction (24 hpi). This later time point allowed for the detection of changes between the advancement of the cycle in control samples vs potential inhibition of the cycle by ectopic expression of the regulatory proteins.

We compared the transcriptome of each sample in triplicate using principal component analysis (PCA). As expected, each set of triplicate biological replicates clustered closely together (Fig. 2A). The Clover control samples clustered in distinct groups depending on isolation timepoint (i.e, 18 vs 24 hpi) (Fig. 2A). For the Euo expressing samples, all the 18 hpi and 24 hpi samples clustered closely together suggesting only small differences in gene expression between the time point samples. This was also seen for HctA expression; the 18 hpi and 24 hpi experimental samples clustered closely together, again suggesting only small differences between time point samples (Fig. 2A). For the HctB 18 hpi and 24 hpi experimental samples each time point clustered separately, but the two clusters were closer to each other than to any of the other experimental conditions. The CtcB 18 hpi and 24 hpi experimental samples were similar, the replicates for each time point clustered tightly together and the 18 hpi and 24 hpi samples clustered closer to each other than to the samples from the other experimental conditions (Fig. 2A). Together, these data suggest that each ectopically expressed protein generated a unique gene expression pattern.

**Figure 2.**
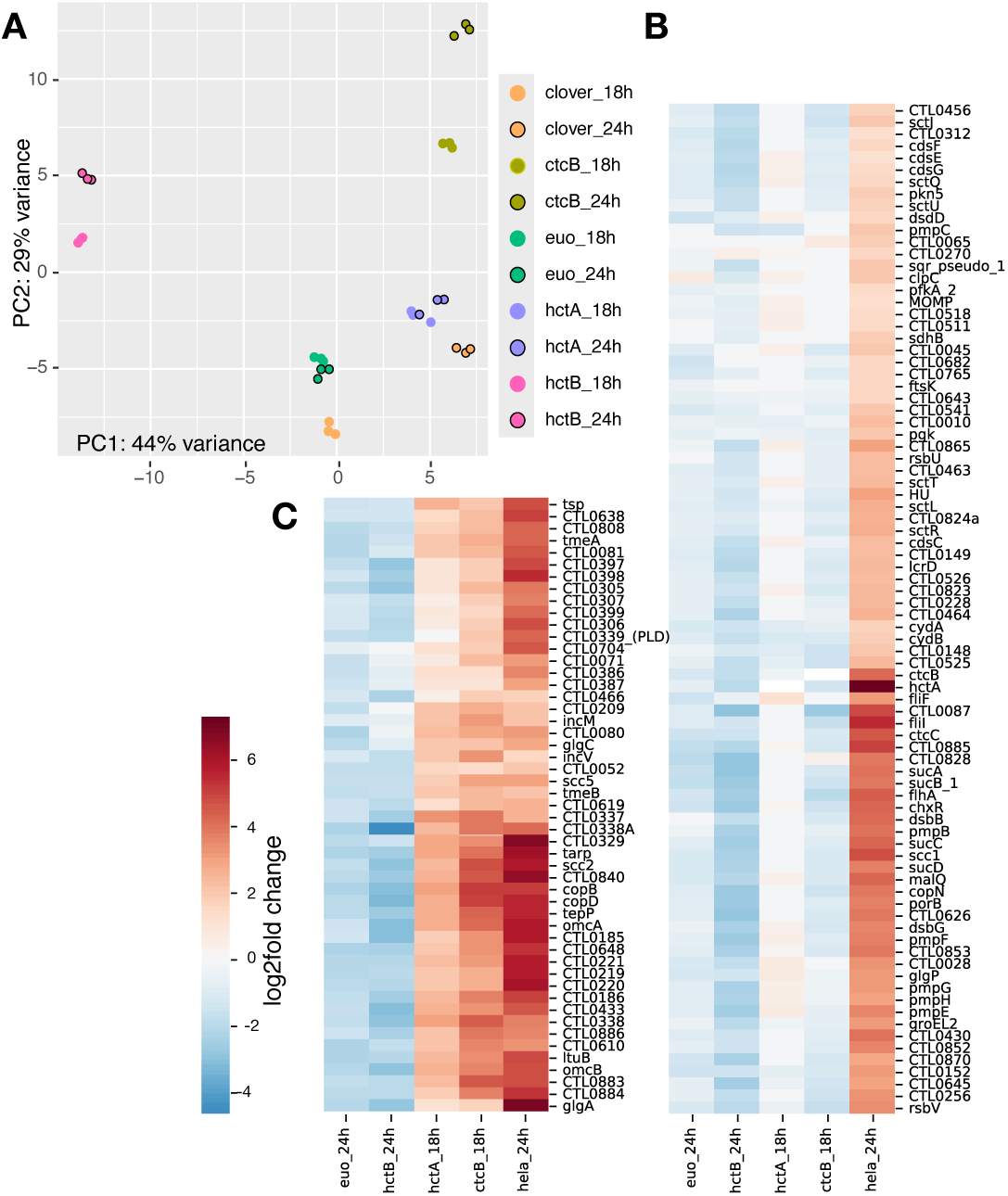
RNA-seq analysis of *Ctr* ectopically expressing Euo, HctA, HctB, or CtcB. (A) For each of the induced samples (n=3) the RNA-seq PCA profiles clustered within the same ectopic expression group but each group had a distinct profile as visualized by plotting the first and second principal component. (B and C) Hierarchically-clustered heatmap plots revealed two distinct late gene regulation groups. (B) The IB cluster group was defined as genes that were upregulated between wt *Ctr* infections at 18 hpi and 24 hpi (late genes) but were not upregulated by the ectopic expression of HctA and CtcB as compared to the Clover control. (C) The EB gene cluster was defined as genes that were upregulated between wt *Ctr* infections at 18 hpi and 24 hpi (late genes) and were induced by ectopic expression of HctA and CtcB as compared to the Clover control.

The RNA-seq data sets were compared to the induced Clover controls; 18 hpi Clover to 18 hpi experimental sample, and 24 hpi Clover to 24 hpi experimental samples. In addition to data from the current analysis, we used the gene expression data from our previously published data set [31] and determined the differential gene expression between *Chlamydia* from the 18 hpi sample and the 24 hpi sample, capturing changes in late gene expression (RB to IB and EB). We compared this differential gene expression pattern to the differential gene expression patterns of the ectopic expression experimental data. We generated a hierarchically-clustered heatmap using the Seaborn clustering algorithm [32] (Fig. 2B and C). The 24 hpi samples for Euo and HctB were used for clustering analysis as these proteins acted as inhibitors and blocked cycle progression (Fig. 2B and C). The 24 hpi samples allowed more time for accumulated changes as the Clover controls progressed to the production of late genes while the Euo and HctB expressing samples did not. The 18 hpi data from the HctA and CtcB ectopic expression experiments were used for cluster analysis as they both acted as inducers (Fig. 2B and C). These changes were most obvious in the 18 hpi samples as the Clover controls had yet to express late genes.

Clustering produced two dominant groups (Fig. 2B and C). Both groups featured genes that were dramatically up-regulated between 18 hpi vs 24 hpi during the wt infection (Fig. 2B and C, HeLa_24h) suggesting that all of these genes would be considered late genes [16,17]. The major difference between the two cluster groups was the changes in gene expression induced by HctA and CtcB ectopic expression (Fig. 2B and C). One of the clusters featured genes that were not dramatically induced upon HctA-FLAG or CtcB-FLAG ectopic expression (Fig. 2B). The other cluster showed the opposite with all the genes upregulated by the ectopic expression of HctA-FLAG or CtcB-FLAG (Fig. 2C). Ectopic expression of Euo-FLAG and HctB-FLAG led to a down-regulation of both sets of genes as compared to the Clover control (Fig. 2B and C). Many of the genes in the first cluster (Fig. 2B) have been shown to be expressed mid-cycle or late-cycle [16,17], while most of the genes in the second-cluster (Fig. 2C) have been identified to be expressed late in the developmental cycle [16,17]. Additionally, many of the second cluster genes have been identified as sigma54/ctcB-ctcC regulated genes [29,33]. These data suggest that the late expressed genes can be divided into two distinct categories; those expressed in the IB (Fig. 2B) and those expressed in the infectious EB (Fig. 2C).

### Gene expression cluster groups map to cell type specific gene expression profiles

Using the clustering data observations, we created a selection criteria to categorize the RNA-seq data into three gene expression groups (Table S1). The first group were genes for which we observed little to no change after Euo ectopic expression when compared to the Clover control and had little to no change in gene expression between 18 hpi and 24 hpi during infection with wt Ctr. We separated the late genes into two groups. The first group was defined as genes whose expression increased between 18 hpi and 24 hpi in the wt infection but were not induced by HctA, CtcB or Euo ectopic expression. The second group was defined as genes whose expression was increased from 18 hpi to 24 hpi in the wt infection and were up-regulated by CtcB and HctA ectopic expression but not increased by Euo ectopic expression. Based on the observation that *euo* was a member of the first group, we defined these genes as RB genes (Table S1). For the two late gene groups we noticed that *hctA*, an IB gene [8,9], was a member of the first group and therefore designated this group as IB genes (Table S1). The second late gene group was designated as EB genes, i.e., the regulon likely involved in the final stage of generating infectious EBs. This group contains the *hctB*, *tarp*, and *scc2* genes, which we have previously shown to be expressed very late in IB to EB developmental progression [8,9].

We next used volcano plots to visualize the individual effects of each of the ectopic expression constructs on changes in gene expression of all *Ctr* genes. The expression changes were plotted (log2fold change) vs statistical significance (-log of the p value) and the RB, IB and EB gene groups listed in Table S1 were highlighted (Fig. 3). We plotted changes in gene expression from 18 hpi to 24 hpi from a wt infection, which as expected indicated that the genes from Table S1 designated as RB genes were largely unchanged in gene expression between 18 hpi and 24 hpi, while both the designated IB and EB genes showed increased expression. RB gene expression was for the most part unchanged when Euo was ectopically expressed, in contrast to dramatic reductions in the expression of both IB and EB designated genes (Fig. 3, Euo 24hpi vs Clover 24hpi). The ectopic expression of both HctA and CtcB dramatically increased the expression of EB genes (Fig. 3, HctA and CtcB 18hpi vs Clover 18hpi). HctB ectopic expression resulted in a dramatic repression in the expression of both IB and EB genes while increasing the relative expression of a subset of RB genes relative to Clover (Fig. 3, HctB 24hpi vs Clover 24hpiE). We noticed that many of the genes that showed increased expression when HctB was ectopically expressed were ribosomal protein genes (Fig. 3, HctB 24hpi vs Clover 24hpi). We interpreted this effect as HctB inhibiting the expression of most genes with the exception of the ribosomal genes, perhaps allowing them to be expressed early during EB to RB differentiation upon the next infection cycle.

**Figure 3.**
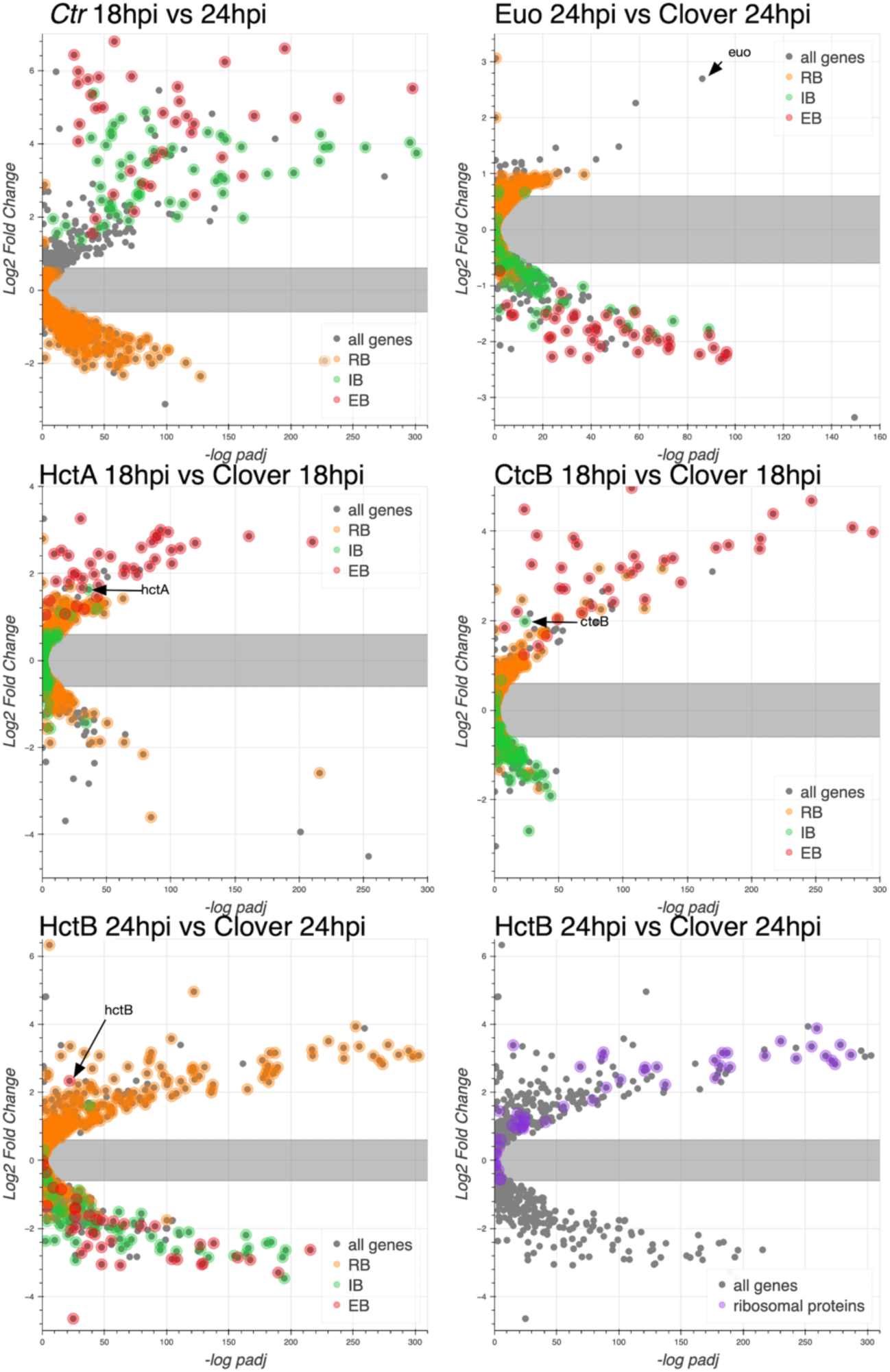
Effects of ectopic expression of Euo, HctA, CtcB and HctB on the gene expression of every *Ctr* gene. RNA-seq differential expression was determined for each gene comparing wt infection at 18 hpi vs 24 hpi, Euo-FLAG expression, HctA-FLAG expression, CtcB-FLAG expression and HctB-FLAG expression vs the control Clover-FLAG. For the wt infection volcano plots show that the expression of RB designated genes (orange) was largely unchanged from 18 hpi to 24 hpi while IB (green) and EB (red) designated genes were dramatically up regulated. For Euo-FLAG expression experiment the RB genes (orange) were largely unchanged while IB genes (green) and EB genes (red) were all down regulated. Ectopic expression of HctA-FLAG resulted in the repression of many of the RB genes (orange) and up regulation of the EB genes (red) but had little impact on the expression of the IB genes (green). CtcB-FLAG expression had very little effect on RB genes (orange) but dramatically upregulated EB genes (red). The ectopic expression of HctB-FLAG resulted in the down regulation of both IB (green) and EB (red) genes but up-regulated many RB genes (orange). Additionally, HctB-FLAG expression resulted in the upregulation of many of the ribosomal protein genes (purple).

### Verification of cell type-specific gene expression by fluorescence *in situ* hybridization

To verify the association of the expression grouped genes with specific cell forms we used fluorescence *in situ* hybridization (FISH) to visualize gene expression in cells expressing GFP and RFP from developmental stage-specific promoters. To this end, we constructed two dual promoter reporter constructs to delineate gene expression from the *euo*, *hctA* and *hctB* promoters which we have shown to be associated with RB, IB and EB cells forms respectively [8,9]. We generated the strains L2-*hctB*prom-mScarlet_*euo*prom-neongreen (L2-BsciEng) and L2-*hctA*prom-mScarlet_*euo*prom-neongreen (L2-AsciEng) which express the RFP mScarlet-I from either the *hctB* promoter or *hctA* promoter along with the GFP protein Neongreen driven by the *euo* promoter. To validate our system, the mRNA expression of *euo*, *hctA* and *hctB* was visualized in each strain using custom FISH probes (Fig. 4). Cells were infected with L2-AsciEng and L2-BsciEng and processed for each FISH probe at 24 hpi. Although dual promoter strains were used in these experiments, the data is presented in a single promoter format to simplify presentation. *Euo* and *hctA* data was processed from L2-AsciEng samples and *hctB* data was processed from L2-BsciEng samples. As expected, *euo* mRNA was observed primarily in the *euo*prom+ (RB) cells and not in either of the *hctA*prom+ (IB) or *hctB*prom+ (EB) cells (Fig. 4A). The *hctA* mRNA was observed in a subset of cells that had overlap with the *hctA*prom signal but not *euo*prom or *hctB*prom signal (Fig. 4B). The *hctB* mRNA was observed in a subset of cells with overlap with *hctA*prom+ and *hctB*prom+ cells but not *euo*prom+ cells (Fig. 4C).

**Figure 4.**
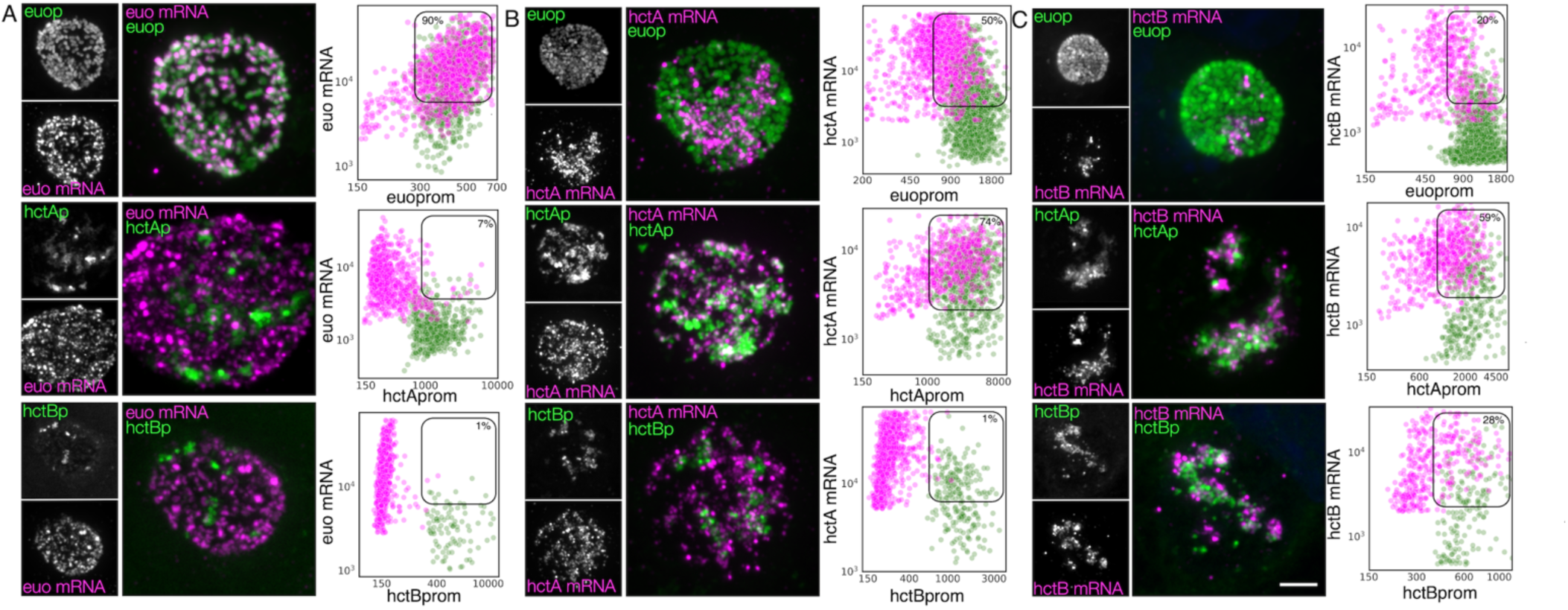
Cell type expression of representative genes from the three gene categories (RB, IB and EB) correspond to the three chlamydial cell forms. Cos-7 cells infected with L2-AsciEng or L2-BsciEng, fixed at 24 hpi and stained using FISH probes for *euo* mRNA, *hctA* mRNA and *hctB* mRNA. (A) Z-projection confocal micrographs showing *euo* mRNA localization in comparison to *euo*prom, *hctA*prom and *hctB*prom activity. Individual chlamydial cells with *euo* mRNA signal from 5 inclusions were identified using TrackMate and the fluorescence intensity for each channel (mRNA and promoter reporter) was plotted (magenta dots). Individual chlamydial cells positive for *euo*prom, *hctA*prom or *hctB*prom signal from 5 inclusions were also identified using TrackMate and their expression intensity for each channel was plotted (green dots). (B) Z-projection confocal micrographs showing *hctA* mRNA localization in comparison to *euo*prom, *hctA*prom and *hctB*prom activity. Individual chlamydial cells with *hctA* mRNA signal from 5 inclusions were identified using TrackMate and the fluorescence intensity for each channel (mRNA and promoter reporter) was plotted (magenta dots). Individual chlamydial cells positive for *euo*prom, *hctA*prom or *hctB*prom signal from 5 inclusions were also identified using TrackMate and their expression intensity for each channel was plotted (green dots). (C) Z-projection confocal micrographs showing *hctB* mRNA localization in comparison to *euo*prom, *hctA*prom and *hctB*prom activity. Individual chlamydial cells with *hctB* mRNA signal were identified from 5 inclusions using TrackMate and the fluorescence intensity for each channel (mRNA and promoter reporter) was plotted (magenta dots). Individual chlamydial cells positive for *euo*prom, *hctA*prom or *hctB*prom signal from 5 inclusions were also identified using TrackMate and their expression intensity for each channel was plotted (green dots). The double positive population was selected (box) and the percentage of the total for the mRNA+ cells (magenta) is indicated in each plot. Size bar = 5µm.

We used the TrackMate plugin in Fiji [34] to identify and quantify both the mRNA signal and promoter reporter signal for each chlamydial cell in five inclusions from each infection. Cells were identified by their promoter reporter signal (green) and also separately by their mRNA fluorescence signal (magenta). The fluorescence intensity was measured and plotted for both channels (FISH and promoter reporter) in both identified populations. Therefore, each plot represents two identified cell populations per reporter, mRNA+ (magenta) population and their corresponding promoter reporter fluorescence intensity, and the promoter reporter+ (green) population and their corresponding FISH signal. This analysis was performed for all three promoter reporters (*euo*p, *hctA*p and *hctB*p) for each FISH mRNA probe (*euo*, *hctA*, and *hctB*) (Fig. 4). The percentage of cells that were single or double positive for each signal was determined and presented in Table S2.

### RB: *euo* FISH

We identified individual chlamydial cells expressing the *euo* mRNA in host cells that were infected with the promoter reporter strains expressing fluorescent proteins from the *euo*prom, *hctA*prom, and *hctB*prom. The analysis indicates that the *euo* mRNA+ cell population (magenta) when plotted for *euo*prom fluorescence and *euo* mRNA fluorescence were primarily double positive (90%) with high levels of both *euo* FISH signal and *euoprom* signal. These *euo* mRNA+ cells from *hctAprom* infections were primarily single positive (93% single+ and 7% double+). This was also observed for the *hctB*prom infections; the *euo* mRNA+ cells when plotted for *euo* mRNA FISH signal intensity against *hctB*prom signal intensity were primarily single positive (99% single+ and 1% double+) (Fig. 4A, Table S2).

We also used TrackMate to identify the promoter reporter positive cell populations (green) and plotted the expression intensities of the promoter reporter signals against the *euo* mRNA FISH signal. The *euo*prom+ cells were predominantly double positive (91%) (high *euo* mRNA signal, high *euo*prom signal), while the *hctA*prom+ cells and *hctB*prom+ cell populations (green) were predominantly single positive (7% and 5% double+ respectively) (low *euo* mRNA signal high *hctA*prom or *hctB*prom signal) (Fig. 4A, Table S2).

### IB: *hctA* FISH

The *hctA* mRNA positive population was identified in the *euo*prom, *hctA*prom and *hctB*prom infected cells using TrackMate (magenta) and the intensities of the *hctA* mRNA FISH signal was plotted against each of the promoter reporter intensity signals. The *hctA* mRNA+ cells from the *euo*prom infection were 50% single positive, likely due to carryover Neongreen protein from RB *euo*prom expression (Fig. 4B, Table S2). The *hctA* mRNA+ cell population (magenta) was mostly double positive (74%) when compared to the *hctA*prom signal. Additionally, the *hctA* mRNA+ population had little to no *hctB*prom signal (1% double positive) (Fig. 4B, Table S2).

We identified the promoter reporter positive chlamydial cells (green) and plotted both the promoter reporter signals and the FISH signal. The *euo*prom+ cell population (green) was mostly single positive (72%) with a small population of double positive cells. The double-positive phenotype was presumably associated with carryover for the long lived Neongreen protein (Fig. 4B, Table S2). The *hctA*prom+ cell population (green) demonstrated a large double positive sub-population (77%) as well as a single positive sub-population that again was likely due to the long half-life of the mScarlet-I protein carried over into the EB population (Fig. 4B, Table S2). The *hctB*prom+ cell population was mostly single positive with little *hctA* mRNA signal (68%) (Fig. 4B, Table S2).

### EB: *hctB* FISH

We performed the same analysis for the *hctB* mRNA+ cells (magenta). The *hctB* mRNA+ chlamydial cells were mostly single positive (80%) when compared to the *euo*prom signal with some detected long-lived Neongreen signal (Fig. 4C, Table S2). The *hctB* mRNA+ cells were generally double positive for the *hctA*prom signal (59%) and double positive for the *hctB*prom signal (28%) (Fig. 4C, Table S2).

For the promoter reporter cells (green), the *euo*prom+ cells had a significant single positive population (82%) and a smaller double positive sub-population, in contrast the *hctA*prom+ cells were both single and double positive for the *hctB* mRNA signal (52% double+ and 48% single+). In the *hctB*prom+ cells there was also both a single and double positive population (61% double+ and 39% single+) (Fig. 4C, Table S2).

The apparent disconnect between mRNA expression profiles and cognate fluorescent protein fluorescence for *euo*, *hctA* and *hctB* FISH results are not unexpected as the fluorescent proteins have a much longer half-life than mRNA. Additionally, fluorescence from mScarlet-I and Neongreen proteins lags mRNA expression as the proteins must be translated and then folded into the mature fluorescent state. Overall, these data indicate that, as expected, *euo* mRNA is expressed in RBs (double positive for euoprom signal and euo mRNA signal), while *hctA* mRNA is expressed in IBs (*hctA*prom+, *hctA* mRNA+ and *hctB*prom negative) and *hctB* mRNA is expressed in late IB/EBs (*hctB*prom+ and *hctB* mRNA+). Therefore, we used this workflow to interrogate cell form gene expression predicted by the RNA-seq clustering, binning and volcano plot analysis.

### Validation of *porB* as an IB gene

Our previous studies showed that the *tarp*, *scc2* and *hctB* promoters were all active much later than the *hctA* promoter [9]. The RNA-seq experiment presented here corroborates this data by placing the corresponding genes in the infectious EB category (Fig. 2C). We also showed that the *hctA* promoter was active in a cell population distinct from the *hctB* promoter making it a likely IB expressed gene [8]. Here we sought to verify an additional gene predicted to be expressed in IBs by the RNA-seq clustering experiment. The porin gene *porB* [35], which clustered with the IB gene group as well as with a proven IB gene *hctA* (Fig. 2B), was selected for this analysis. Cells were infected with either L2-AsciEng or L2-BsciEng, fixed at 24 hpi and probed for the *porB* mRNA. Confocal images were taken and viewed as z projections (Fig. 5A). The *porB* mRNA signal (magenta) did not completely overlap with the *euo*prom+ signal (green) and appeared to be expressed in a subset of cells (Fig. 5A). The *porB* mRNA had significant but not complete overlap with the *hctA*prom+ cells (green) and almost no overlap with the *hctB*prom+ cells (Fig. 5A). Using the TrackMate protocol described above we identified the promoter reporter expressing populations, *euo*prom, *hctA*prom, and *hctB*prom (green) and then separately, the *porB* mRNA+ population (magenta) and measured the fluorescence intensity of each channel (fluorescent reporter proteins and mRNA signal within each population). The *porB* mRNA+ population (magenta) in the *euo*prom channel experiment were a mix of single and double positive cells (70% double+ and 30% single+) (Fig. 5A and Table S2). This was also true for the *euo*prom+ population (green) (73% double+ and 27% single+). In comparison, the *porB* mRNA+ population (magenta) in the *hctA*prom channel experiment were also double positive and single positive (56% double+ and 44% single+). Additionally, the *hctA*prom+ population (green) was primarily double positive cells (78%) (Fig. 5A and Table S2). In the *hctB*prom channel experiment, the *porB* mRNA signal+ population (magenta) were primarily single positive (97%) and did not have appreciable *hctB*prom fluorescence. Conversely the *hctB*prom+ population (green) was primarily single positive 80% with low *porB* mRNA signal. Taken together, the *porB* mRNA expression pattern was similar to that of the *hctA* mRNA expression pattern strongly suggesting *porB* is expressed primarily in the IB cell form.

**Figure 5.**
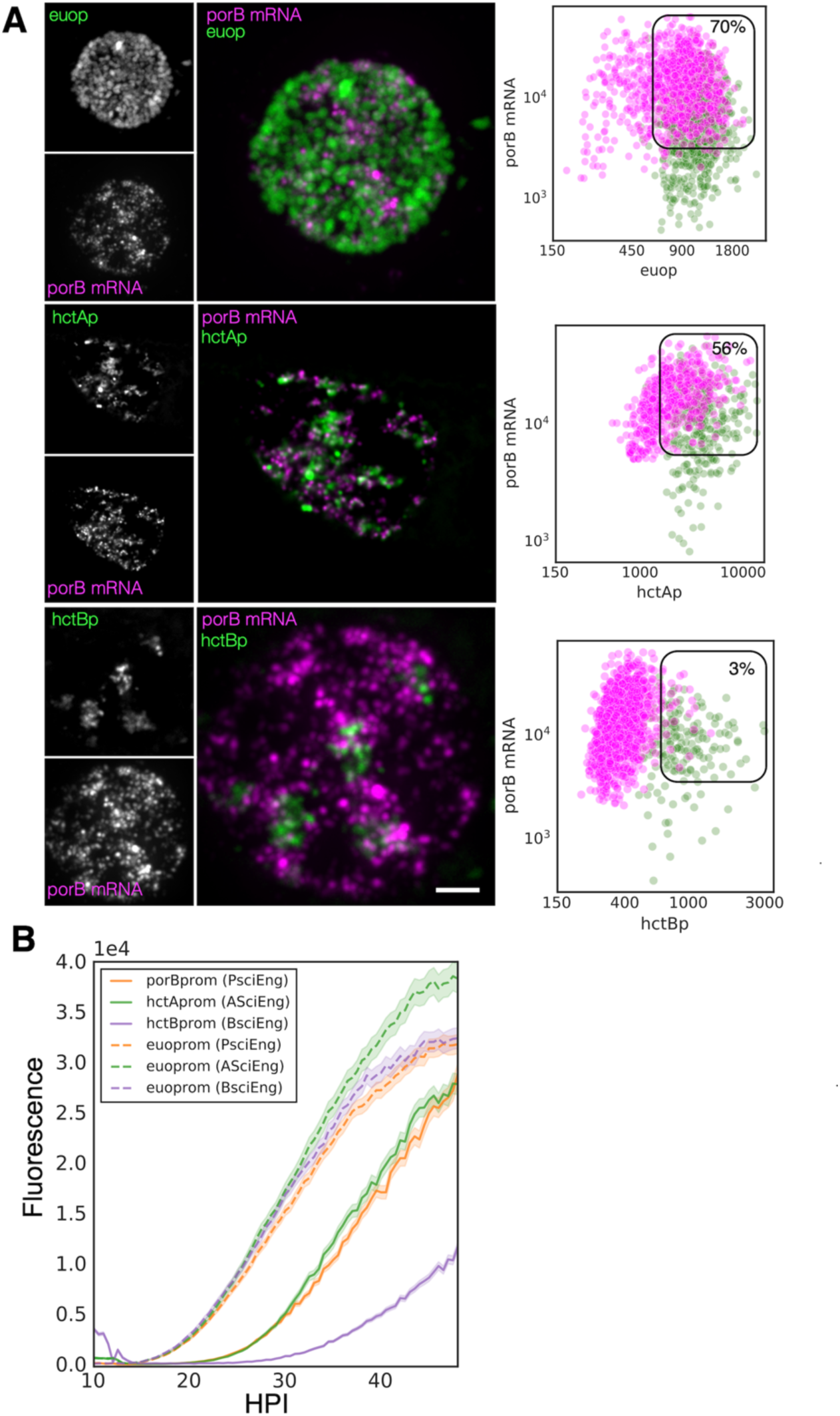
PorB gene expression is consistent with being classified as an IB gene. (A) FISH analysis of *porB* mRNA expression in comparison to *euo*prom, *hctA*prom and *hctB*prom activity at 24 hpi. Confocal micrographs of L2-AsciEng and L2-BsciEng infected cells probed for *porB* mRNA expression from 5 inclusions using Molecular instruments FISH probes. TrackMate was used to identify the *porB* mRNA+ cell population and measure the FISH fluorescent signal as well as the *euop*rom, *hctA*prom and *hctB*prom fluorescent signal. The intensity for both channels for each cell was plotted (magenta dots). The *euo*prom, *hctA*prom and *hctB*prom+ cell population were also identified using TrackMate and the signal from the FISH channel and fluorescent protein channels were plotted on the same graphs (green dots). The double positive population was selected (box) and the percentage of the total for the mRNA+ cells (magenta) is indicated in each plot. (B) Cells were infected with L2-PsciEng and the developmental gene expression kinetics were compared to those of L2-AsciEng, and L2-BsciEng. The *euo*prom expression kinetics were comparable for all strains with expression first detected at ∼15 hpi. *PorB*prom expression kinetics were nearly identical to *hctA*prom expression kinetics first detected at ∼20 hpi while *hctB*prom expression was initiated at ∼26 hpi. Error cloud for fluorescent reporters represents SEM. n > 20 inclusions per strain.

We next evaluated the kinetics of the activity of the *porB* promoter to determine if the kinetics were similar to the *hctA* promoter [8,9]. The *hctA* promoter of AsciEng was replaced with the promoter region of *porB* (-137bp to +30bp) and transformed into *Ctr* to create L2-PsciEng. We used live cell imaging to measure the expression of Neongreen driven by *euo*prom and mScarlet-I driven by *porB*prom. Cells were infected with PsciEng at an MOI ∼0.3 and imaged for both the Neongreen and mScarlet-I fluorescence at 10 hpi every 30 minutes for a further 48 hours. For comparisons, L2-AsciEng and L2-BsciEng strains were imaged in parallel as we have previously shown that *euo* promoter activity is detected at ∼15 hpi followed by the *hctA* promoter and finally the *hctB* promoter [8,9]. The kinetics of the *porB* promoter mirrored that of the *hctA* promoter. *Euo*prom activity was detected at ∼15 hpi followed by the activity of *porB*prom and *hctA*prom at ∼20 hpi and by *hctB*prom activity starting at ∼26 hpi (Fig. 5B).

Cell form-specific promoter activity was also evaluated in the L2-PsciEng strain (Fig. S2). Cells were infected with L2-PsciEng, fixed at 16 hpi and 24 hpi, and evaluated using confocal microscopy. At 16 hpi the inclusion contained primarily *euo*prom+ cells forms (bright green) and little to no *porB*prom signal. The inclusions at 24 hpi contained both an *euo*prom+ subset of chlamydial cells as well as a subset of cells that were *porB*prom+ (Fig. S2). Taken together these data suggest that *porB*, as predicted by the RNA-seq clustering data, can be considered an IB gene.

### Predicted cell type expression of T3SS genes

We noticed an intriguing expression pattern of the T3SS structural genes in the gene expression profile data that suggested cell form specific expression. To explore this observation, we plotted the effects of ectopic expression of the regulatory proteins on the T3SS operons. We used our wt RNA-seq data [36], operon prediction software [37] and the RT-PCR data published by Hefty et al. [38] to annotate the T3SS operons (Table S3) and plotted the expression data using volcano plots. These plots revealed that the majority of the T3SS operons were regulated in an IB-like pattern of gene expression, i.e. up-regulated between 18 hpi and 24 hpi, repressed by Euo and HctB ectopic expression, but not induced by CtcB or HctA ectopic expression (Fig. 6). The exception to this pattern were the two operons for the T3SS translocons; (*CTL0238, lcrH, copB_2* and *copD_2,* (*CTL0238-* op)) and (*scc2, CTL0840, copB* and *copD,* (*scc2*-op)). The four genes in the *CTL0238*-op were regulated like RB genes while the four genes in the *scc2*-op were regulated like EB genes (Fig. 6).

**Figure 6.**
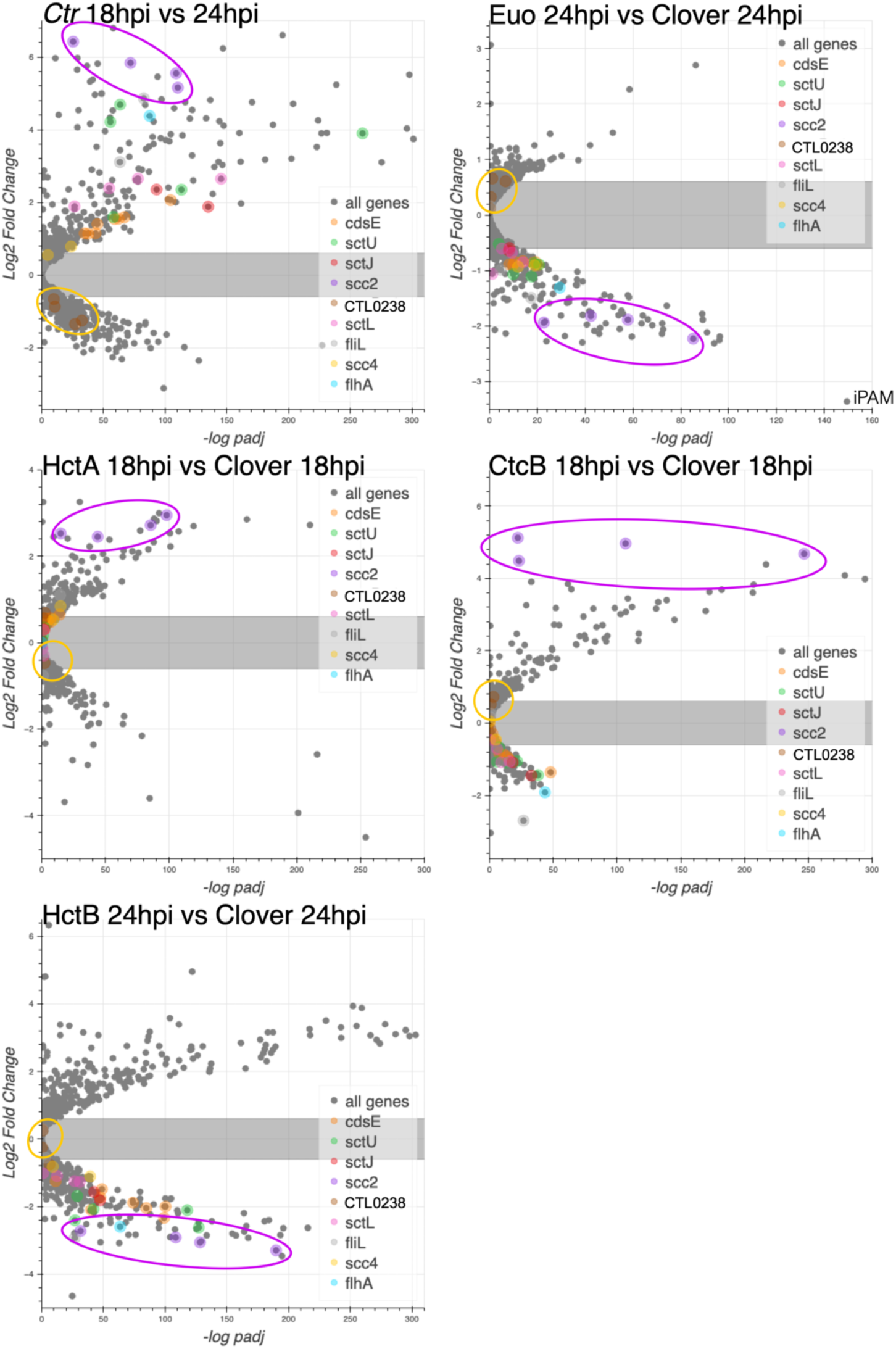
The effects of ectopic expression of Euo, HctA, CtcB and HctB on T3SS structural genes. The log2fold change RNA-seq differential expression data from the ectopic expression experiments were plotted against the -log of the p value (-log padj) and the operons for the T3SS were highlighted. For the wt *Ctr* 18 hpi vs 24 hpi samples most of the T3SS structural genes were upregulated while for the Euo ectopic expression experiment most of these operons were downregulated. Again, like the IB genes in the HctA and CtcB ectopic expression experiments, most of the structural genes were downregulated or unchanged. Two operons did not follow this pattern, the *CTL0238*-op and *scc2*-op. Both operons encode the components of the T3SS translocon. The four genes in the *CTL023*8-op (gold circle) were regulated like RB genes while the four genes in the *scc2*-op (purple circle) were regulated like EB genes.

### Validation of cell type expression of T3SS structural operons by FISH

We next investigated the cell type expression of two of the T3SS operons predicted to be expressed in the IB, the *sctU* operon and the *sctJ* operon. The *sctU* operon (*sctU*-op) encodes the genes *sctU*, *sctV*, *lcrD*, *copN*, *scc1*, and *malQ*, while the *sctJ* operon (*sctJ*-op) includes the genes *sctJ*, *sctK*, *sctL*, *sctR*, *sctS*, and *sctT*. We used custom FISH probes for *sctU* through *lcrD* for the detection of the *sctU-op* mRNA and *sctL* to *sctR* for the detection of *sctJ*-op mRNA. Cell monolayers were infected with L2-AsciEng and L2-BsciEng at an moi ∼0.3 and processed for FISH staining at 16 and 24 hpi (Fig. 7 *sct*Jo and Fig. S3 *sct*Uo). FISH signal was not observed in the RB cells (*euo*prom+) at 16 hpi for either *sctU*-op (Fig. S3A) or *sctJ*-op (Fig. 7A). In the infections fixed at 24 hpi, the FISH staining for both operons was observed in cells distinct from *euo*prom+ and *hct*Bprom+ cells (Fig. 7B, *sctJ*o and Fig. S3B, *sctU*o). However, both T3SS operon mRNAs were detected in a subset of the *hctA*prom+ cell population (Fig. 7B, *sctJ*o and Fig. S3B, *sctU*o). We again used our TrackMate workflow to quantitate these data. Cells were identified by their promoter reporter signal (green) and separately by their mRNA/FISH signal (magenta). The fluorescence intensity was measured and plotted for both channels (FISH (magenta) and promoter reporter (green)) in both identified populations as described for Fig. 4. As *sctJ*-op and *sctU*-op results were very similar, only the sctJ-op data analysis is discussed in detail below.

**Figure 7.**
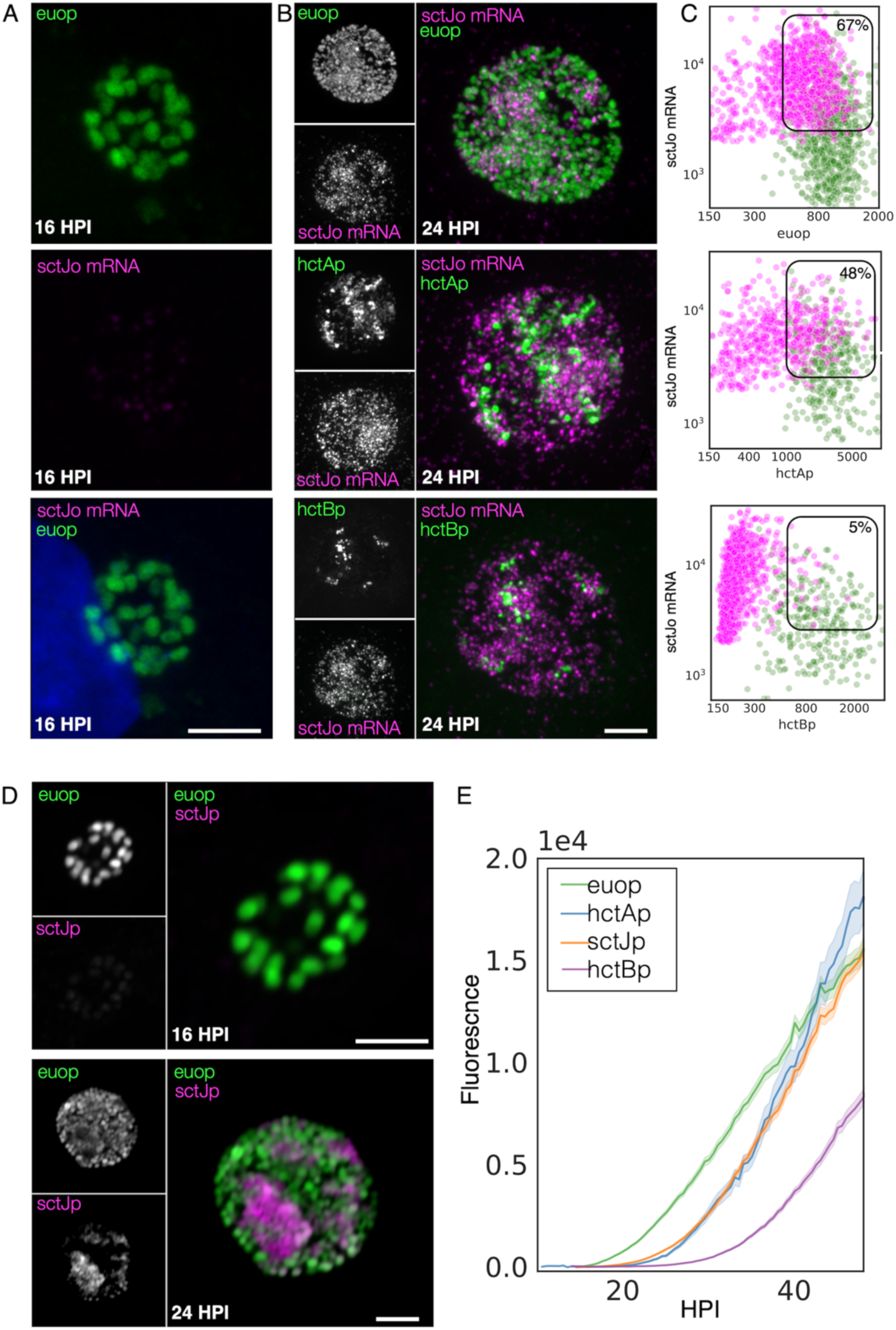
IB cell type expression of the T3SS structural operon *SctJ-*op. (A) Cells were infected with L2-AsciEng for 16 hpi and fixed and stained using a FISH probe (*sctL* to *sctR*) to the mRNA for the T3SS structural operon *sctJ*-op, the RB control *euo* and the IB control *hctA*. All cells were positive for *euo*prom expression (green). The FISH stained cells were only positive for *euo* mRNA (magenta) and were negative for *hctA* mRNA (magenta) and *sctJ-op* mRNA (magenta). (B) Cells were infected with L2-AsciEng and L2-BsciEng for 24 hpi and fixed and stained using FISH for the *sctJ*-op mRNA. For the *euo*prom sample, the *sctJ*-op FISH signal (magenta) was present in a distinct subset of cells and not in the majority of the *euo*prom+ cells (green). (C) TrackMate was used to identify the *sctJ*-op mRNA+ cells from 5 inclusions and the signal for *euo*prom and FISH were quantified for each *sctJ*-op+ cell and plotted (magenta dots). The converse was also performed, the *euo*prom+ cells were identified and the *euo*prom signal and FISH signal was quantified for each *euo*prom+ cell and plotted (green dots). The FISH signal was also compared to the *hctA*prom expression pattern and showed subsets of cells that were stained for both *sctJ*-op mRNA and *hctA*prom expression as well as non overlapping populations. The *sctJ*-op mRNA+ cells were again identified using TrackMate and the signal for *hctA*prom and FISH were quantified for each *sctJ*-op+ cell and plotted (magenta dots). Each *hctA*prom+ cell was also identified and the FISH and *hctA*prom signal was determined and plotted (green dots). The *sctJ-*op FISH staining was also compared to the expression from the *hctB*prom reporter. The *sctJ*-op mRNA FISH staining was again present in a subset of cells but showed little overlap with the *hctB*prom fluorescent signal. The FISH signal and *hctB*prom signal were measured in both cell populations (*sctJo* mRNA+ cells and *hctB*prom+ cells) and plotted, *sctJo* mRNA+ cells magenta dots and *hctB*prom+ cells green dots. Both populations were primary single positive, either *sctJ*-op mRNA high or *hctB*rpom high but rarely both. The double positive population for mRNA+ cells was selected (box) and the percentage of the total is indicated. (D) Cos-7 cells infected with L2-JsciEng (*sctJ* promoter driving scarlet-I) were fixed at 16 hpi and 24 hpi and imaged. The 16 hpi inclusions contain primarily *euo*prom expressing cells (green) with little *sctJ*prom scarlet-I signal. At 24 hpi there are two dominant cell populations euoprom+ and sctJprom+ cells. Size bar = 5µm. (E) The kinetics of *sctJ*prom activity was determined and compared to that of the *euo*prom, *hctA*prom and *hctB*prom. Cos-7 cells infected with L2-JsciEng, L2-AsciEng and L2-BsciEng and imaged every 30 minutes starting at 10 hpi until 48 hpi. The *euo*prom signal began to increase at ∼15 hpi while the *sctJ*prom and *hctA*prom signal began to increase at ∼22 hpi followed by the *hctB*prom activity at 28 hpi.

### *sctJ*-op: Expression in RB cells

We identified the mRNA+ cell population (magenta) and quantified both the *euo*prom signal intensity and the FISH signal intensity. For the *sctJ*-op mRNA+ cells, there was both a double positive population (high mRNA signal and high *euo*prom signal) and single positive population (67% double and 33% single+). We also quantified the mRNA expression in RBs by identifying the *euo*prom+ cells (green) and measuring the *sctJ*-op FISH signal and plotted this against the *euo*prom signal intensity (Fig. 7C *euop,* Table S2). The *euo*prom+ population (green) was both single and double positive for both operons (56% double+ and 44% single+) (Fig. 7C *euo*p, Table S2). At 16 hpi there was no measurable *sctJ* mRNA signal in any of the cells.

### *sctJ*-op: Expression in IB cells

The *sctJ*-op mRNA+ cell population (magenta) was identified and both the *hctA*prom signal intensity and the FISH signal intensity was quantified and plotted. For the *sctJ*-op mRNA+ cells (magenta), there was both a double positive population and a single positive population (48% and 52% respectively) (Fig. 7C *hctAp,* Table S3). For the *hctA*prom+ cell population (green) there was both a single (*hctA*prom) and double positive (*hctA*prom and mRNA) population (65% and 35% respectively) (Fig. 7C *hctA*p, Table S2). These data further suggest that the *sctU*-op and *sctJ*-op were expressed in the IB cell type. It’s likely that the *hctA*prom+ single positive population are late IB/EB cell forms that are becoming EBs and have repressed *sctJ*-op expression.

### *sctJ*-op: Expression in EB cells

In contrast, the mRNA+ cell population for the *sctJ*-op in the *hctB*prom expressing cells were distinct single positive (mRNA signal) populations (5% double+ and 95% single+) (Fig. 7C *hctB*p, Table S2). Additionally, the *hctB*prom+ cell population was also primarily single positive (*hctB*prom). These data suggest that the *sctJ*-op was not expressed in the EB cell forms. Combined, these overall expression patterns of the *sctJ* operon were very similar to that of the *hctA* mRNA and *porB* mRNA FISH, supporting an IB-like gene expression pattern.

To determine cell type specificity for expression of the *sctJ* operon, we replaced the *hctA* promoter in the AsciEng construct with the *sctJ* promoter (120 bp upstream of the ATG start of *sctJ*) and transformed it into *Ctr* L2 creating L2-JsciEng. Cells were infected with L2-JsciEng, fixed at 16 hpi and 24 hpi and cell form specificity was evaluated using confocal microscopy. At 16 hpi only the Neongreen signal was detected (Fig. 7D). There were two obvious cell populations present at 24 hpi, one brightly expressing the Neongreen protein from the *euo* promoter and a second population that was brightly expressing the mScarlet-I protein from the *sctJ* promoter (Fig 7D). In addition to confocal microscopy, we used live cell imaging to measure the kinetics of expression of Neongreen driven by the *euo* promoter and mScarlet-I driven by the *sctJ* promoter. Cells were infected with L2-JsciEng at an MOI ∼0.3 and imaged for both Neongreen and mScarlet-I fluorescence every 30 minutes from 10 hpi for 55 hours. For comparisons, L2-AsciEng and L2-BsciEng strains were imaged in parallel [8,9]. The kinetics of the *sctJ*prom activity mirrored that of *hctA*prom (Fig. 7E). Overall, this data supports the observation that the *sctJ* and *sctU* operons are expressed primarily in the IB cell form.

### FISH-based analysis of cell type expression of the T3SS translocon operons

As mentioned above, the *Ctr* genome encodes two operons for the T3SS translocon each of which contain four genes, the *CTL0238*-op and the *scc2*-op [39]. This duplication is conserved in all the vertebrate-infecting chlamydial species. The expression profiles from our clustering data and volcano plots suggested that the two translocon operons are expressed in different cell types; the *CTL0238*-op in RBs and the *scc2*-op in EBs (Fig. 6). To verify differential cell type expression, host cells were infected with L2-BsciEng, fixed at 24 hpi and probed with custom FISH probes designed against C*TL0238*-op and *scc2*-op. Confocal micrographs showed that the mRNA FISH signal for *CTL0238*-op heavily overlapped with the *euo*prom channel but was distinct from *hctB*prom+ cells (Fig. 8A, *CTL0238*-op mRNA). In contrast, the mRNA signal for the *scc2*-op was distinct from the *euo*prom+ cells but almost completely overlapped the *hctB*prom+ cells (Fig. 8B, *scc2*-op mRNA).

**Figure 8.**
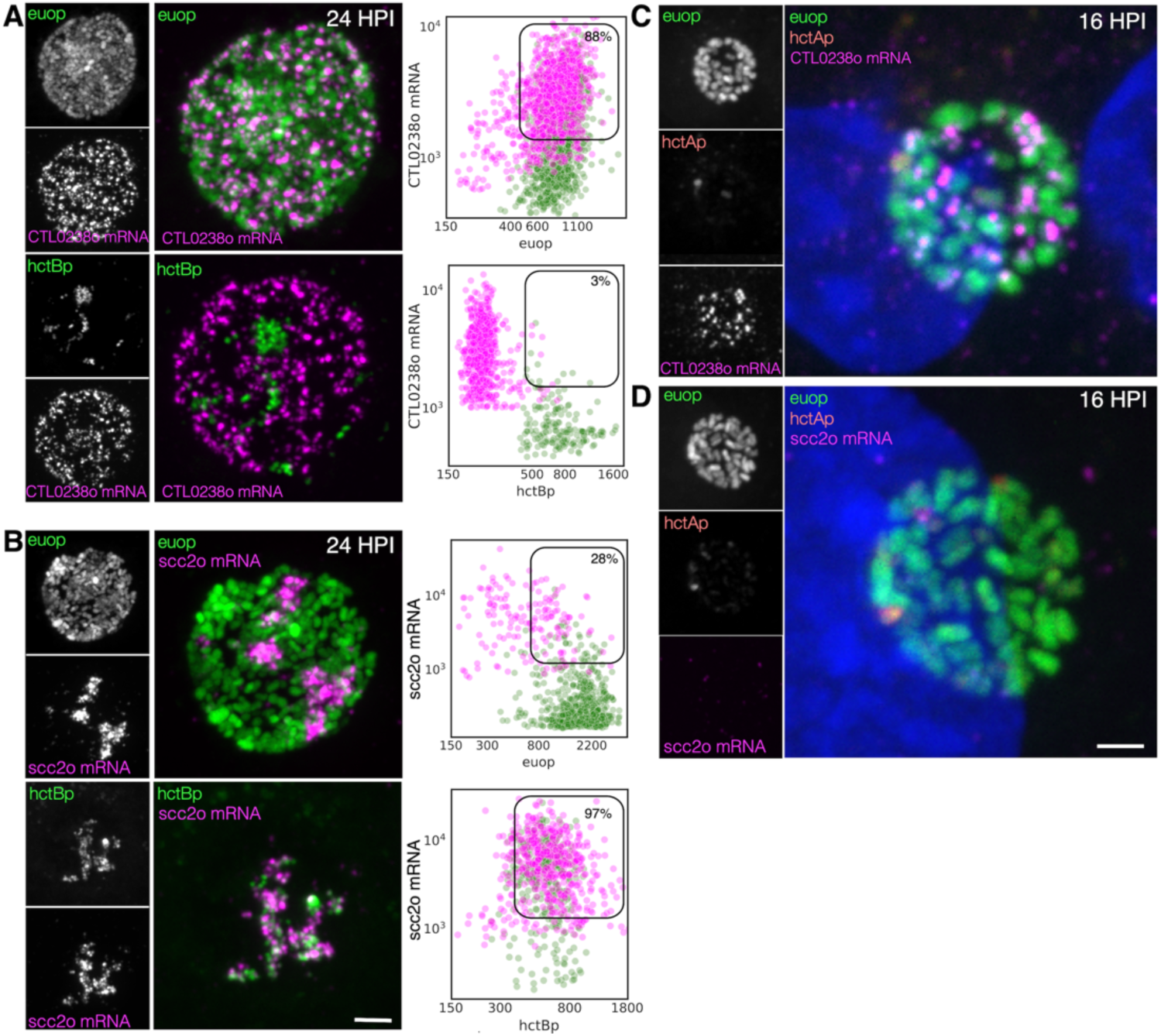
Cell type expression of the two T3SS translocons. A) Cos-7 cells were infected with L2-BsciEng for 24 hpi and stained for the mRNA expression of the *CTL0238*-op using FISH (magenta), *euo*prom expression (green) and *hctB*prom expression (green). Individual chlamydial cells with *CTL0238*-op mRNA signal from 5 separate inclusions were identified using TrackMate and the fluorescence intensity for each channel (mRNA and promoter reporter) was plotted (magenta dots). Individual chlamydial cells positive for *euo*prom, or *hctB*prom signal from 5 separate inclusions were also identified using TrackMate and the expression intensity for each channel (mRNA and promoter reporter) was plotted (green dots). B) Cos-7 cells were infected with L2-BsciEng for 24 hpi and stained for the mRNA expression of the *scc2*-op using FISH. *Scc2*-op FISH signal in magenta, *euo*prom and *hctB*prom signal in green. Individual chlamydial cells positive for *scc2*-op mRNA signal from 5 inclusions were identified using TrackMate and the fluorescence intensity for each channel (mRNA and promoter reporter) was plotted (magenta dots). Individual chlamydial cells positive for *euo*prom, or *hctB*prom signal from 5 inclusions were also identified using TrackMate and the expression intensity for each channel (mRNA and promoter reporter) was plotted (green dots). The double positive population for mRNA+ cells was selected (box) and the percentage of the total is indicated. (C) Host cells infected with AsciEng and fixed at 16 hpi were probed for *CTL0238*-op mRNA and *scc2*-op mRNA (D) expression using FISH. *Euo*prom expression (green) had significant overlap with *CTL0238-*op mRNA signal. For the *scc2*-op the FISH signal was undetected. Size bar = 5µm.

We again quantified this expression pattern using our TrackMate workflow. Chlamydial cells were identified by their promoter reporter signal (green) and then separately by their mRNA fluorescence signal (magenta). The fluorescence intensity was measured and plotted for both channels; FISH (magenta) and promoter reporter (green).

### *CTL0238*-op: Expression in RB cells

We identified the *CTL0238*-op mRNA+ cell population and quantified both the *euo*prom signal intensity and the FISH signal intensity. For the *CTL0238*-op mRNA+ cells, there was primarily a double positive population (88%) (high mRNA signal and high *euo*prom signal) (Fig. 8A, Table S2). We also quantified the mRNA expression in RB cells (*euo*prom+ cells). The *CTL0238*-op FISH signal was plotted against the *euo*prom signal intensity and the *euo*prom+ cells were mostly double positive (71%) with an additional single positive population (29%) (Fig. 8A, Table S2). This single positive (*euo*prom+, *CTL0238*-op mRNA-) population is likely due to the long halflife of the GFP protein.

### *CTL0238*-op: Expression in EB cells

In contrast, the *CTL0238*-op mRNA+ cell population when plotted for mRNA signal and the *hctB*prom signal was a distinct single positive population (97%) (*CTL0238*-op mRNA+, hctBprom-) (Fig. 8A, Table S2). We also identified the *hctB*prom+ cell population and plotted the mRNA signal and *hctB*prom signal. This population was also primarily single positive (89% (*hctB*prom+, *CTL0238*-op mRNA-) (Table S2).

### scc2-op

The *scc2-op* mRNA FISH quantification showed the opposite results (Fig 8B). The *scc2-op*+ mRNA cells were primarily single positive when plotted against the *euo*prom signal (28%) and double positive when plotted against the *hctB*prom signal (97%) (Fig. 8B, Table S2). The *euo*prom+ cell population was only 10% double positive while the *hctB*prom+ cells were primarily double positive (89%) for *scc2-op*+ mRNA (Fig. 8B, Table S2).

To further highlight the differential expression of *CTL0238*-op mRNA and *scc2*-op mRNA, we infected cells with L2 AsciEng and processed the samples for FISH at 16 hpi when most of the chlamydial cells are RBs. As expected, at 16 hpi essentially all the cells were green RBs (*euo*prom+) with little to no red IB (*hctA*prom+) cells. The *euo*prom+ cells were all positive for the *CTL0238*-op FISH signal (Fig. 8C, *CTL0238*-op). In contrast, the *scc2*-op FISH signal was undetectable in *euo*prom+ cells at 16 hpi (Fig. 8D, *scc2*-op).

Taken together, these data support the observation that the two translocon operons are differentially regulated and are expressed in distinct cell forms. The *scc2*-op is expressed in late IB/EB cells while the *CTL0238*-op is expressed in RB cells.

### Predicted cell type expression of T3SS effectors

In general, T3SS translocons are involved in interacting with host membranes to facilitate the secretion of T3SS effectors into target cells [40]. During infection, *Ctr* secretes effectors into/through two membrane systems, the host cell plasma membrane and, once inside the cell, the chlamydial inclusion membrane. Additionally, the chlamydial T3SS is known to secrete different kinds of effectors, soluble proteins as well as the integral membrane inclusion (Inc) proteins [41–44]. Using volcano plots we asked how the genes encoding the soluble effector proteins (Table S4) and Inc proteins (Table S5) were regulated by the ectopic expression of Euo, HctA, HctB and CtcB. The vast majority of the soluble T3SS effector genes were regulated like EB genes; higher in 18-24 hpi and induced by HctA and CtcB ectopic expression but down regulated by Euo and HctB ectopic expression (Fig. 9A). In contrast, most of the *incs* were expressed as RB genes except for *incM* and *incV,* which were expressed like EB genes (Fig. 9B).

**Figure 9.**
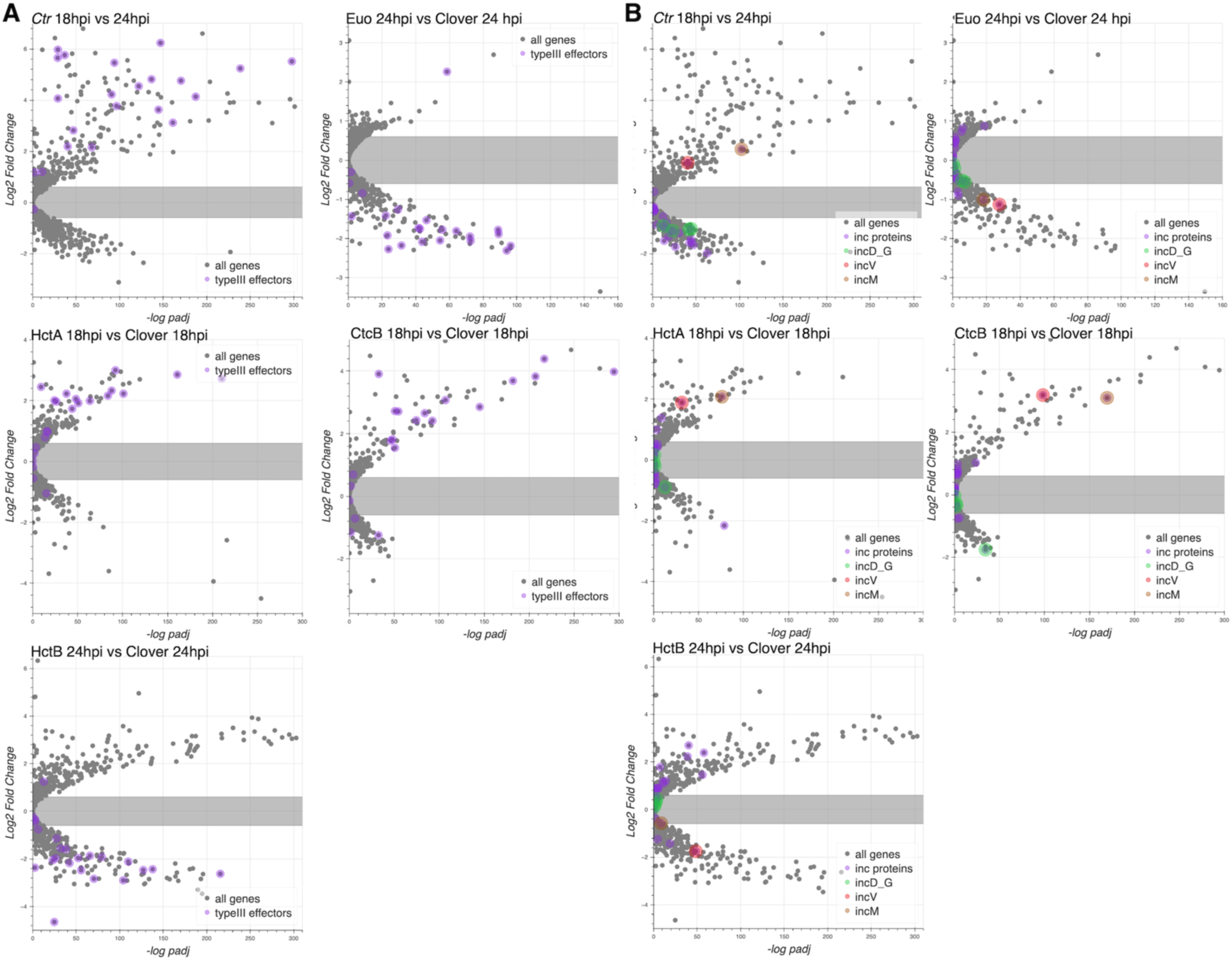
Effects of ectopic expression of Euo, HctA, CtcB and HctB on the expression of T3SS effectors. RNA-seq differential expression data (log2fold change) plotted vs the -log of the P value (-log padJ) for *Ctr* ectopically expressing Euo, HctA, CtcB and HctB. (A) The T3SS effectors are highlighted in purple. (B) All the *inc* protein genes are highlighted in purple while the genes for the *incD-G* operon are highlighted in green, *incV* and *incM* are highlighted in red and orange respectively.

### Cell type expression of *inc* genes by FISH

The volcano plots suggested that the majority of the *inc* effector genes were expressed in RBs. However, two *inc* genes (*incV* and *incM)* stood out as potential EB genes (Fig. 9B). To determine if the putative late Incs, *incV* and *incM,* were expressed late in RBs or were bona fide EB genes, we compared mRNA expression of a known RB expressed Inc, *incD,* to the expression of *incV* and *incM* using FISH. Cells infected with L2-BsciEng were probed for the expression of *incD*, *incV* and *incM* mRNA at 16 hpi (mostly RBs) and 24 hpi (all three cell forms). Confocal microscopy revealed that, as expected, *incD* was expressed in *euo*prom+ RB cells at both 16 hpi and 24 hpi and not in *hctB*prom+ EBs present at 24 hpi (Fig. 10A). Conversely, *incV* and *incM* mRNA could not be detected at 16 hpi (no EBs) and were expressed exclusively in *hctB*prom+ EBs (Fig. 10B, *incV* and Fig. S4, *incM*).

**Figure 10.**
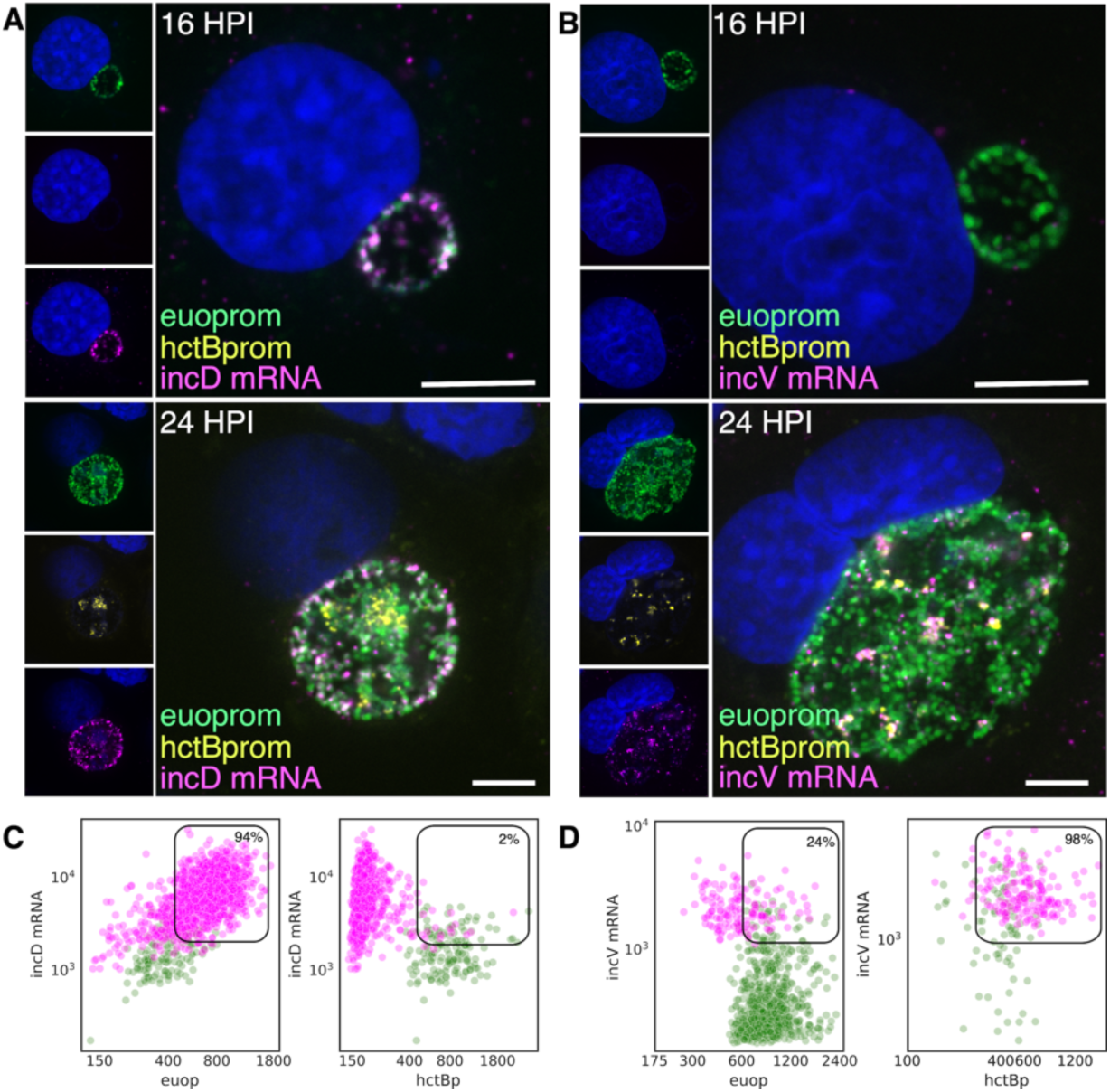
Cell type expression of *incD* and *incV*. Cos-7 cells infected with L2-BsciEng for 16 and 24 hpi and stained for *incD* and *incV* mRNA expression using custom FISH probes. (A) The *incD* mRNA (magenta) was visibly expressed in the *euo*prom+ (green) RB cells at 16 hpi while *hctB*prom signal was not detected. At 24 hpi the *incD* mRNA signal (magenta) overlapped with the *euo*prom signal (green) but was separate from the *hctB*prom+ cells (yellow). (B) The *incV* mRNA signal (magenta) was undetected at 16 hpi. At 24 hpi the *incV* mRNA signal showed overlap with the *hctB*prom signal (yellow) but not the *euo*prom signal (green). C) Individual chlamydial cells positive for *incD* mRNA signal were identified from 5 separate inclusions at 24 hpi using TrackMate and the fluorescence intensity for each channel (mRNA and promoter reporter) was plotted (magenta dots). Individual chlamydial cells positive for *euo*prom, or *hctB*prom signal were also identified using TrackMate and the expression intensity for each channel (mRNA and promoter reporter) was plotted (green dots). D) Individual chlamydial cells positive for *incV* mRNA signal from 5 separate inclusions at 24 hpi were identified using TrackMate and the fluorescence intensity for each channel (mRNA and promoter reporter) was plotted (magenta dots). Individual chlamydial cells positive for *euo*prom or *hctB*prom signal were also identified using TrackMate and the expression intensity for each channel (mRNA and promoter reporter) was plotted (green dots). The double positive population for the mRNA+ cells was selected (box) and the percentage of the total is indicated. Size bar = 5µm.

We next quantified expression of *incD* and *incV* in RBs and EBs from inclusions from the 24 hpi experiments using our TrackMate workflow. Chlamydial cells were identified by their promoter reporter signal (green) and separately by their mRNA fluorescence signal (magenta) and the signal for both populations was plotted.

### *incD*: Expression in RB cells

We identified the *incD* mRNA+ cell population (magenta) and plotted both the *euo*prom signal intensity and the FISH signal intensity (Fig. 10C, Table S2). For the incD mRNA+ cells there was primarily a double positive population (94%) (high *incD* mRNA signal and high euoprom signal). We also quantified the mRNA expression in RB cells (euoprom+ cells). The *incD* FISH signal was plotted against the *euo*prom signal intensity and the *euo*prom+ cells were mostly double positive (90%).

### *incD*: Expression in EB cells

In contrast, the *incD* mRNA+ cell population when plotted for mRNA signal and the *hctB*prom signal was a distinct single positive population (98%) (*incD* mRNA+, *hctB*prom-) (Fig. 10C, Table S2). We also identified the *hctB*prom+ cell population and plotted the mRNA signal and *hctB*prom signal. This population was primarily single positive (75%) (*hctB*prom+, *incD* mRNA-).

### incV

We only analyzed and plotted the *incV* data as *incV* and *incM* showed similar FISH results. The *incV*+ cells were primarily single positive (76%) when plotted against the *euo*prom signal and double positive when plotted against the *hctB*prom signal (98%) (Fig. 10D, Table S2). The *euo*prom+ cell population was also primarily single positive (97%) and the *hct*Bprom+ cells were primarily double positive (60%) (Fig. 10D, Table S2). These data support the hypothesis that *incD* is indeed an RB gene and that *incV* and *incM* are EB genes.

## Discussion

The chlamydial developmental cycle has traditionally been defined by the timeline of the infection. The infectious EB invades the host cell and differentiates into the RB cell form that then begins to divide. The genes involved in this process have been described as the early genes. After EB to RB differentiation RBs replicate and the gene expression associated with this timeframe is usually considered the chlamydial midcycle. Genes upregulated from ∼24 hpi until cell lysis, when EBs accumulate in the inclusion, are considered late genes [16–18]. We have dissected the developmental cycle and developed a model based on cell type transitions [8]. Our model suggests that the developmental cycle is best described by a programmed cell production model [8]. In this model, the EB enters the host cell (through the use of premade effectors) and initiates immediate early protein synthesis (EB to RB differentiation genes) to begin the EB to RB differentiation process. The EB to RB differentiation process takes ∼ 10 hours to complete. The completion of EB to RB differentiation is defined by the first division of the nascent cell resulting in RB cells. At this stage the RBs expand in number through cell division, amplifying the infection. Our model suggests the RBs mature during this amplification stage, ultimately producing daughter cells with asymmetric fates. One daughter cell becomes the IB cell form while the other remains an RB. Our model defines the IB as the cell type committed to EB formation. The mature RBs at this stage continue to replicate producing one IB and one RB. The IBs never re-enter the cell cycle and instead transition into the infectious EB which takes ∼ 10 hours to complete [8].

In this study, we ectopically expressed four transcriptional regulatory proteins that all blocked the progression of the developmental cycle. The effects of expression of these regulatory proteins using RNA-seq was determined and compared using a clustering algorithm which resulted in three distinct regulation patterns. The first cluster contained genes that were unaffected by the ectopic expression of Euo and were not upregulated between 18 hpi and 24 hpi of a *Ctr* L2 wt infection. The second cluster consisted of genes whose expression increased from 18 hpi to 24 hpi of a wt infection but were not induced by ectopic expression of HctA or CtcB. The third cluster of genes were upregulated between 18 hpi and 24 hpi and by the ectopic expression of both HctA and CtcB. These groups fit well into the major cell categories in our model; RBs, IBs and EBs. Using the clustering observation, we created selection criteria based on changes in gene expression from our RNA-seq experiments. We were able to categorize 639 of 902 genes (70%) into one of the RB, EB, or IB categories. The genes that could not be assigned were either expressed at levels too low to have confidence in the expression pattern or had a unique expression pattern that did not fit into the three categories suggesting potential unique roles in chlamydial biology. This study focused on determining gene expression through measuring mRNA and it remains to be determined if any of these genes are translationally regulated as well.

The RB cell is the replicating cell form leading to expansion of cell numbers. Based on the changes in gene expression after ectopic expression of Euo, HctA, CtcB or HctB we found that 532 genes were regulated as RB genes. This category included cell replication genes, genes involved in protein synthesis, genes for many of the Inc proteins and *euo*. Based on our selection criteria this group likely encompasses both potential constitutive genes (expressed in RBs, IBs and potentially early EBs) as well as RB-specific genes such as *euo*, *incD* and the *CTL0238*-op which we show were expressed only in the RB cell form.

The IB cell type is the transitional form between the RB and the EB and is currently poorly defined. We define the IB cell type as the committed step to EB formation; the IB is the cell form that exits the cell cycle and begins the program to transition into the infectious EB [8,22]. Our data identified 67 genes that are likely expressed specifically in the IB cell type. The functions of these genes vary widely. We identified two porin genes (*porB* and *CLT0626*), two disulfide isomerases (*CTL0149* and *CTL0152*) and six polymorphic outer membrane proteins (*pmpB*, *C, E*, *F*, *G*, and *H*) as IB genes, suggesting dramatic changes to the outer membrane of the IB as it transitions into the EB.

The EB cell is the infectious cell form that is “terminally” differentiated. Once formed in the inclusion, the EB maintains an infectious phenotype through active metabolism but has very low levels of protein expression [36]. Here, we define the EB regulon as the genes expressed during the late IB to EB maturation phase. Of the 46 EB genes, 18 had been previously shown to be directly regulated by the sigma54 alternative sigma factor and 4 were reported to be sigma28 regulated genes [29,33]. The regulation of the remaining 24 genes is unknown. As both HctA ectopic expression and the ectopic expression of CtcB induce the expression of the EB genes, the EB regulon is likely regulated by a complex shift in gene expression and activation of the sigma54 and sigma28 regulons is a part of this shift.

We tested one of the predicted IB genes, *porB*, and showed that its regulation, both by promoter specific gene expression in chlamydial cells and by its developmental kinetics, matched that of the IB gene *hctA.* This was further confirmed using FISH to demonstrate cell type gene expression matched that of *hctA*. We have previously published the kinetics of the *euo*, *hctA* and *hctB* promoters and showed that the promoter activities fit into the RB, IB and EB model [8,9]. Here we combined these promoter reporter strains with FISH and demonstrated that the *euo* mRNA was expressed primarily in RBs, that *hctA* mRNA was expressed in IBs, and that *hctB* mRNA was expressed in EBs demonstrating the usefulness of FISH for identifying cell type specific gene expression.

Overall, these data support a model that includes (at least) three dominant cell forms: the RB, the IB and the EB. These cells have dramatically different gene expression profiles and phenotypes. The EB has been well characterized as it is the infectious form, does not replicate and has a dramatically condensed nucleoid. The nucleoid structure is due in part to the binding of the two histone like proteins, HctA and HctB to the chromosome [23,26]. Our data indicates that the construction of the compact nucleoid occurs in two distinct and temporally separated steps [8,9]. HctA is expressed as an IB gene and, when ectopically expressed, resulted in the expression of the EB genes suggesting that HctA expression is an important regulator of the IB to EB transition. HctB on the other hand is expressed as an EB gene and, when ectopically expressed, resulted in the inhibition of the expression of most genes with the exception of the ribosomal protein genes. An intriguing hypothesis is that the ribosomal protein genes are potentially free of inhibition in the mature EB which could in turn allow protein synthesis to be rapidly reinitiated upon infection to aid in EB to RB differentiation, without a requirement for complete removal of HctA and HctB from the chromosome. Taken together, these data suggest that the transition from the IB to EB occurs in two steps; 1) HctA chromosomal binding potentially turns off RB and IB genes, allowing EB genes to become expressed, and 2) HctB is expressed late in EB formation creating the final condensed nucleoid and turning off the majority of gene expression but potentially sparing the ribosomal genes.

Volcano plots of the effects of ectopic expression of the four regulatory genes support the categorization of most chlamydial genes into the RB, IB and EB categories. We specifically focused on the expression of the T3SS operons and observed that the majority of the operons for the structural components were IB-like in their regulation. This was verified using FISH for both the *sctJ* operon (*sctJ*, *sctK*, *sctL, sctR, sctS, and sctT*) and the *sctU* operon (*sctU, sctV, lcrD, copN, scc1, and malQ*). Additionally, the promoter for the *sctJ* operon was active in the IB cell form. While the majority of the T3SS structural operons were expressed as IB genes, the two translocon operons (*CTL0238, lcrH, copB_2, copD_2*) and (s*cc2, CTL0840, copB, copD*) were predicted by clustering and volcano plots to be expressed in RB and EB cells respectively. This prediction was again verified by FISH in the context of dual promoter reporter strains.

The observation that the two translocons were expressed in distinct cell forms (*CTL0238*-op in RBs and *scc2*-op in EBs) prompted us to determine the expression of the T3SS effectors. *Ctr* encodes two classes of effectors, soluble and inclusion membrane embedded proteins (Incs) [42,45,46]. The data from this study showed that the majority of the Inc protein effectors (28 out of 36) were expressed as RB genes while the majority of the soluble T3SS effectors (17 out of 23) were expressed as EB genes and that none of the soluble effectors were expressed as RB genes. This pattern supports an intriguing model; the *scc2*-op translocon translocates soluble effectors as the EB contacts host cells and mediates entry events, while the *CTL0238*-op is expressed early during the EB to RB differentiation process in the nascent inclusion and translocates the transmembrane Inc effectors. Whether this separation is temporal or whether the two translocons are specialized for the translocation of soluble vs. inclusion membrane effectors is currently unknown. Interestingly, although the majority of the Inc proteins were expressed as RB genes there were two Incs (*incV* and *incM*) that were determined to be expressed in EBs. In addition to their regulation pattern, we also verified that *incV* and *incM* were EB genes using FISH. Both *IncV* and *IncM* are involved in the establishment of early inclusion functions and are expressed late in the developmental cycle [16,47,48]. We hypothesize that these “pre-loaded” Inc proteins are among the first to be secreted from internalized *Ctr* after the *CTL0238*-op is deployed.

*Ctr* communicates and reprograms the host cell to create and maintain its intracellular replication niche in part through the use of the T3SS. We were surprised that the majority of the T3SS operons for the structural components of the system were expressed as IB genes. This expression pattern along with the cell type specific expression of the translocons (one in the RB and one in the EB) and effectors suggests that the T3SS is constructed, deployed and secretes effectors in a cell type-specific manner that is likely a critical component of the complex developmental cycle and host cell reprogramming.

Our model depicted in Figure 11 suggests that the EB binds to and enters cells in part through the deployment of soluble effectors and the *scc2*-op translocon expressed during EB development. After entry, EB to RB differentiation begins and the RB genes are expressed, this includes the *CTL0238*-op translocon which deploys the Inc proteins for the creation of the inclusion replication niche and the genes required for chlamydial replication leading to RB amplification. After an amplification period the RB matures into a stem cell-like cell form and begins to produce IBs [8]. The T3SS structural components are assembled in the IB and this facilitates maturation to the EB form [8,9]. That the IB and not the RB expresses the genes for the construction of the T3SS suggests that the T3SS apparatus deployed on the EB cells remains on the RBs and is diluted with every round of replication. It is unclear if the secretion system is partitioned equally or is retained in a subset of RBs. Intriguingly, this supports a proposed role of T3SS dilution in cell form maturation/development put forth previously [43,49,50].

**Figure 11.**
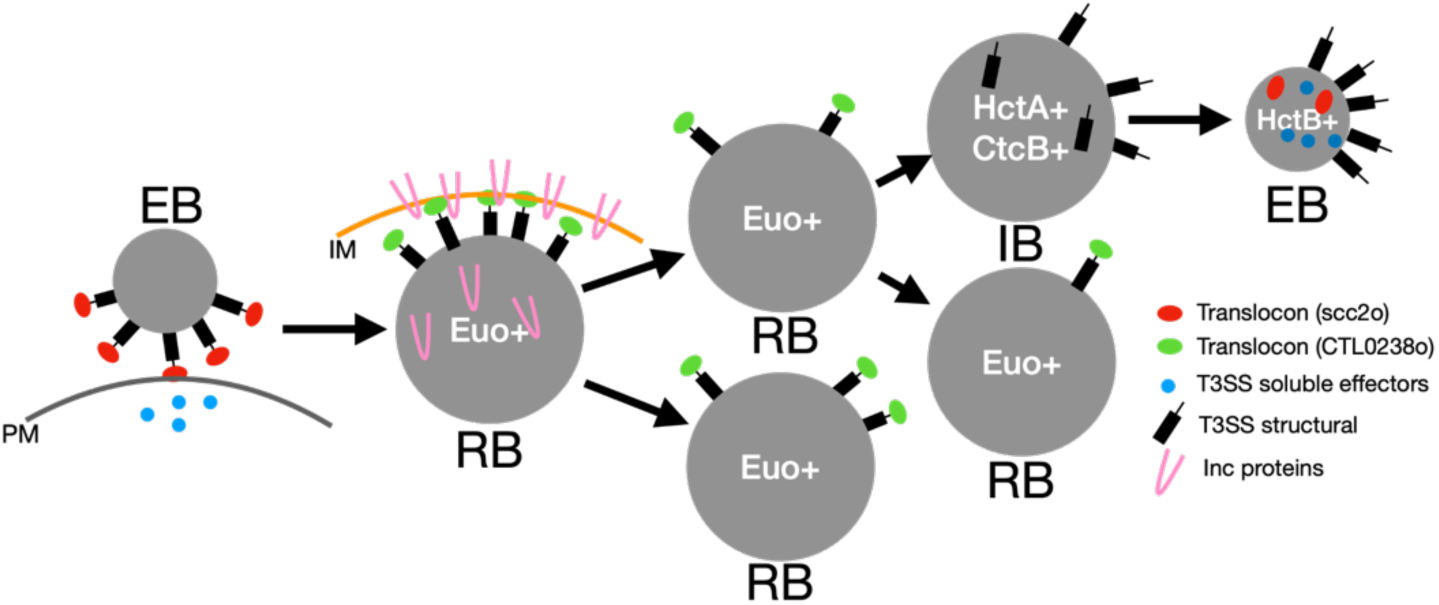
Model of cell type specific deployment of the T3SS. In this model the *scc2-*op translocon secretes effectors across the plasma membrane (PM) for host cell entry. The *scc2*-op translocon is replaced in the RB with the *CTL0239-*op translocon for the secretion of the Inc proteins across the inclusion membrane (IM). The structural components of the T3SS are then reconstructed during the IB to EB maturation phase.

The IB also expresses the histone-like DNA binding protein, HctA. Previous studies have shown that when expressed in *E. coli*, HctA can alter gene expression in a gene specific manner [23]. Our data suggest that HctA has an important role in shifting gene expression from the IB pattern to the EB genes. This is likely in conjunction with the CtcB/C two component regulatory system and sigma54 [29,33]. The EB genes, as previously mentioned, include the majority of the soluble T3SS effectors and the *scc2*-op translocon as well as the HctB DNA binding protein. We hypothesize that EB gene expression loads the EB with the invasion-related proteins and HctB shuts down the majority of gene expression creating the final condensed nucleoid, the final step of EB formation. This prepares the EB for the initiation of the next round of infection (Fig. 11).

DNA replication is tightly controlled during the *Ctr* developmental cycle; only the RB cell form replicates the chromosome and the IB and EB cells contain a single fully replicated chromosome [22,51]. The role of the control of DNA replication in regulating gene expression is currently unknown. However, it is intriguing to speculate that DNA replication could contribute to changes in DNA supercoiling which has been shown to play a role in gene expression during the chlamydial developmental cycle [52–54].

Our data has highlighted three categories of gene expression that define the three major phenotypic cell forms, the RB, IB and EB. However, future studies are needed to define the regulatory circuits and DNA elements that create these cell form-specific expression patterns. Identification of cell type gene expression of a large percentage of the chlamydial genome will aid the determination of the functions of the many hypothetical genes encoded in the chlamydial genome. Understanding the function of many of these genes has been hampered by the mixed cell environment of the chlamydial inclusion. Additionally, with the emerging genetic tools available to investigate the functional roles of genes during infection, knowing in which cell type a gene is expressed will improve the interpretation of the data.

## Materials and Methods

### Cell Culture

Cell lines were obtained from the American Type Culture Collection. Cos-7 cells (CRL-1651) were grown in RPMI-1640, supplemented with 10% FBS and 10 μg/mL gentamicin (Cellgro). *Chlamydia trachomatis* serovar L2 (LGV Bu434) was grown in Cos-7 cells. Elementary Bodies (EBs) were purified by density gradient (DG) centrifugation essentially as described [55] following 48 h of infection. EBs were stored at −80°C in Sucrose Phosphate Glutamate (SPG) buffer (10 mM sodium phosphate [8mM K2HPO4, 2mM KH2PO4], 220 mM sucrose, 0.50 mM l-glutamic acid, pH 7.4) until use.

### Vector Construction

All constructs used p2TK2-SW2 [56] as the backbone and cloning was performed using the In-fusion HD EcoDry Cloning kit (FisherScientific). Primers and geneblocks (gBlocks) were ordered from Integrated DNA Technologies (IDT) and are noted in Table S6. For the ectopic expression of Clover, Euo, CtcB and HctA the T5 promoter (*E. coli* sigma70 constitutive promoter) and the E riboswitch was used for conditional translational expression control using the inducer, theophylline (Tph) [30]. For the ectopic expression of HctB the Tet promoter was used in conjunction with the E riboswich to confer both transcriptional and translational expression control (Tet-JE-hctB) and has been described previously [30]. The *hctA, hctB*, *euo* and *ctcB* ORFs were amplified from *Ctr* L2(434) using the primers indicated in Table S6.

To create the Scarlet-I reporters *hctB*prom_Scarlet-*euo*prom_neongreen (BsciEng), *hctA*prom_Scarlet-*euo*prom_neongreen (AsciEng), *porB*prom_Scarlet-*euo*prom_neongreen (PsciEng) and *sctJ*prom_Scarlet-*euo*prom_neongreen (JsciEng) the gBlock mScartlet-I (Table S6) was cloned into BMELVA [8] to replace the mKate RFP gene. The degradation tag LVA was then removed from neongreen using the primers indicated. The *hctA*, *porB* and *sctJ* promoters were amplified and used to replace the *hctB* promoter using the primers indicated to create AsciEng, PsciEng and JsciEng respectively.

### Chlamydial Transformation and Isolation

Transformation of Ctr L2 was performed essentially as previously described [9]. Briefly, 1×10^8^ EBs + >2µg DNA/well were used to infect a 6 well plate. Transformants were selected over successive passages with 1U/ml penicillin G or 500µg/ml spectinomycin as appropriate for each plasmid. The new strain was clonally isolated via successive rounds of inclusion isolation (MOI, <1) using a micromanipulator. Clonality of each strain was confirmed by isolating the plasmid, transforming into *E. coli* and sequencing six transformants.

The chlamydial strains L2-E-euo-FLAG, L2-E-hctA-FLAG, and L2-E-ctcB-FLAG were induced at the indicated times with 0.5 mM Tph. As described previously, *Ctr* could not successfully be transformed with the E-hctB-FLAG construct, therefore we developed a tet-riboJ-E promoter system that combines both transcriptional and translational control to hctB-FLAG expression, creating the strain L2-tet-J-E-hctB-FLAG [30]. Expression of HctB-FLAG was induced with 0.5 mM Tph+30ng/ml anhydrotetracycline (aTc).

### Replating Assay

*Ctr* were isolated by scraping the infected monolayer into media and pelleting at 17200 rcfs. The EB pellets were resuspended in RPMI via sonication and seeded onto fresh monolayers in a 96-well microplate in a 2-fold dilution series. Infected plates were incubated for 24 hours prior to fixation with methanol and stained with 4′,6-diamidino-2-phenylindole (DAPI) and *Ctr* MOMP Polyclonal Antibody, FITC (Fishersci). The DAPI stain was used for automated microscope focus and visualization of host-cell nuclei, and the anti-*Ctr* antibody was used to visualize the *Ctr* to identify and count inclusions. Inclusions were imaged using a Nikon Eclipse TE300 inverted microscope utilizing a scopeLED lamp at 470nm and 390nm, and BrightLine band pass emissions filters at 514/30nm and 434/17nm. Image acquisition was performed using an Andor Zyla sCMOS in conjunction with μManager software. Images were analyzed using ImageJ software and custom scripts. Statistical comparisons between treatments were performed using an ANOVA test followed by Tukey’s Honest Significant Difference test.

### Transmission Electron Microscopy

For analysis of the structure of *Ctr* upon ectopic protein expression, cell monolayers were infected with the indicated strain at an moi of 0.5 and induced with 0.5mM Tph at 15 hpi. Infected cells were released from the plate with Trypsin-EDTA at 30 hpi, rinsed with 1xPBS and the pellet was fixed with EM fixative (2%PFA, 2% Glutaraldehyde, 0.1M Phosphate Buffer, pH 7.2) overnight at 4°C. Fixed pellets were rinsed and dehydrated before embedding with Spurr’s resin and cross sectioned with an ultramicrotome (Riechert Ultracut R; Leica). Ultra-thin sections were placed on formvar coated slot grids and stained with uranyl acetate and Reynolds lead citrate. TEM imaging was conducted with a Tecnai G2 transmission electron microscope (FEI Company; Hillsboro, OR).

### RNA-Seq

Expression of each protein was induced at 15 hpi with either 0.5 mM Tph and 30ng/ml anhydrotetracycline (HctB) or 0.5 mM Tph (Clover, HctA, CtcB and Euo) and the *Ctr* isolated at 18 and 24 hpi on ice. Total RNA was isolated from the indicated infections and treatments. Briefly, the infected monolayer was scraped into ice cold PBS, lysed using a Dounce homogenizer and the *Ctr* isolated over a 30% MD-76R pad. Total RNA was isolated using TRIzol reagent (Life Technologies) following the protocol provided and genomic DNA removed (TURBO DNA-free Kit, Invitrogen). Both prokaryotic and eukaryotic rRNAs were depleted using Illumina Ribo-Zero Plus. The enriched RNA samples were quantified and the libraries built and barcoded by the IBEST Genomics Resources Core at the University of Idaho. The libraries were sequenced by University of Oregon sequencing core using the Illumina NovaSeq platform. The chlamydial reads were analyzed by aligning to the published *Ctr* L2 Bu 434 genome using the Bowtie2 aligner software [57]. The aligned chlamydial reads were quantified for each chlamydial ORF using HTseq. For each sample ∼1X10^6^ read pairs were counted for 904 chlamydial orfs resulting in about 1000X coverage for each orf. Statistical analysis and normalization of read counts was accomplished using DESeq2 in R [58]. Log2fold change and statistics were also calculated using DESeq2. Heatmaps and hierarchical clustering were generated and visualized using Python with Pandas and the Seaborn visualization package [32]. The raw reads and HT-seq counts are accessible from the NCBI’s Gene Expression Omnibus with the accession number of GSE287626. Volcano plots were constructed from the log2fold change data using Python and the Bokeh plotting library (Bokeh Development Team).

### RNA fluorescence in situ hybridization (RNA-FISH)

All FISH probes were designed by Molecular Instruments (Los Angeles, CA) using the sequence indicated in (Table S7). Cos7 monolayers seeded on coverslips were infected with the indicated strains at an MOI ∼ 0.3. Infected cells were fixed at the indicated times in 4% paraformaldehyde (PFA) for 10 min at RT at 24 hpi, washed 2x with 1XPBS and dehydrated overnight at −20°C in 70% EtOH. Samples were probed and the signal was amplified as described by the protocol provided by Molecular Instruments with the exception that DAPI was added to the final wash to visualize DNA. Coverslips were mounted on a microscope slide with MOWIOL® mounting solution (100 mg/mL MOWIOL® 4-88, 25% glycerol, 0.1 M Tris pH 8.5).

Fluorescence images were acquired using a Nikon spinning disk confocal system with a 60x oil-immersion objective, equipped with an Andor Ixon EMCCD camera, under the control of the Nikon elements software. Images were processed using the image analysis software ImageJ (http://rsb.info.nih.gov/ij/). Representative confocal micrographs displayed in the figures are maximal intensity projections of the 3D data sets, unless otherwise noted.

### Live cell imaging

Monolayers of Cos7 cells were grown in a glass bottom 24-well plates and infected with the promoter reporter strains L2-BsciEng, L2-AsciEng and L2-PsciEng. Live cell imaging of the developing inclusions was started at 8 hpi using an automated Nikon epifluorescent microscope equipped with an Okolab (http://www.oko-lab.com/live-cell-imaging) temperature-controlled stage and an Andor Zyla sCMOS camera (http://www.andor.com). Multiple fields of view from each well were imaged every fifteen minutes. The fluorescence intensity of each inclusion over time was tracked using the ImageJ plugin TrackMate [34] and the results were averaged and plotted using python and matplotlib [59].

## Acknowledgements

We would like to thank Dr. Dan Rockey for careful reading and editing of the manuscript.

## Supplemental tables

**Table S1**. Chlamydial genes designated as RB, IB, EB expressed genes and their regulation from ectopic expression of Euo, HctA, CtcB and HctB

**Table S2**. Quantification of single chlamydial expression plots.

**Table S3**. T3SS structural operons.

**Table S4**. T3SS effector gene regulation from ectopic expression of Euo, HctA, CtcB and HctB.

**Table S5**. Inc gene regulation from ectopic expression of Euo, HctA, CtcB and HctB.

**Table S6**. List of primers used to construct plasmids

**Table S7**. List of FISH probes and the location on the *Ctr* genome

## Supplemental figures

**Fig. S1.**
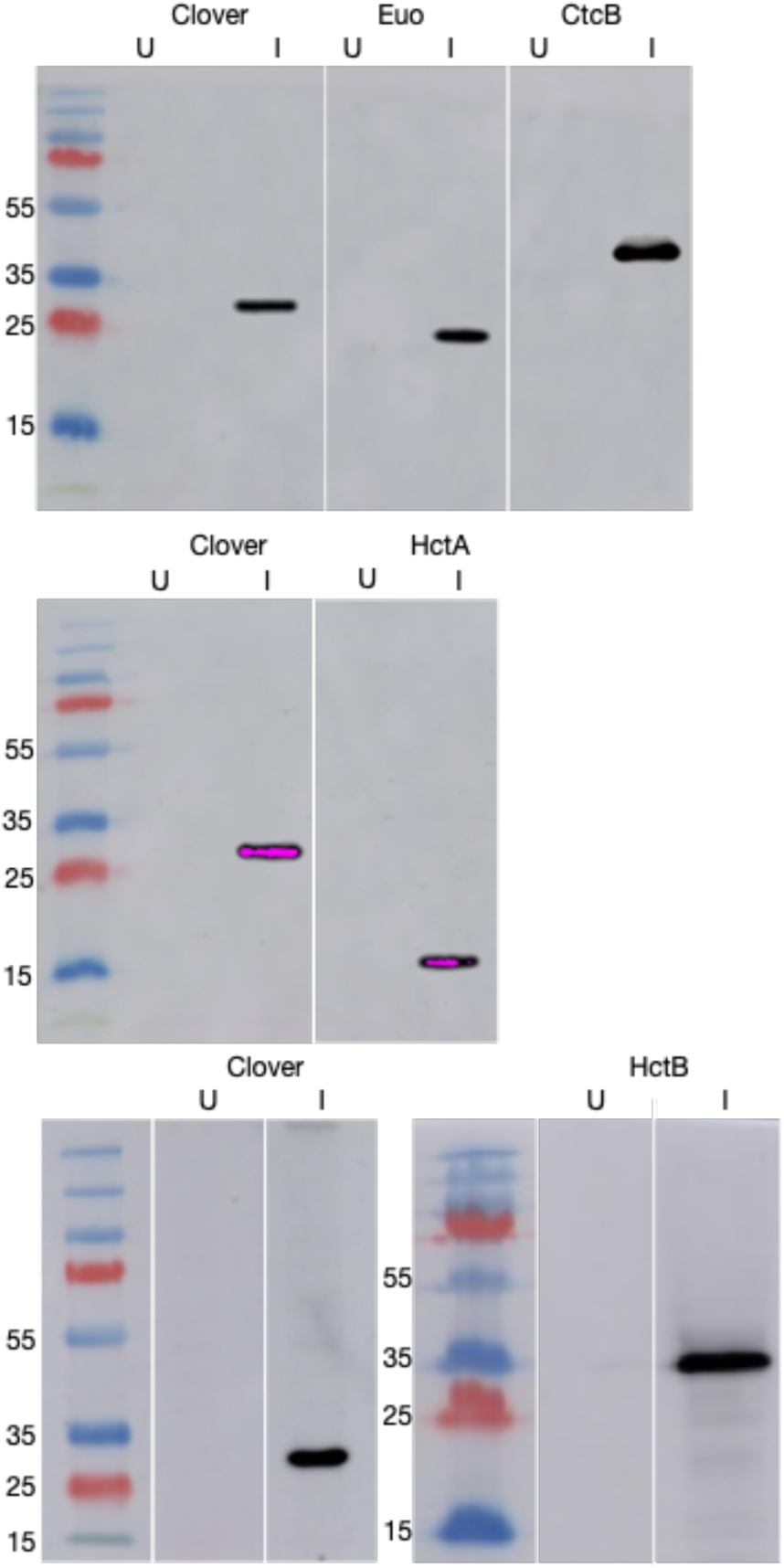
Western analysis of ectopically expressed Clover, Euo, HctA, CtcB and HctB. To ensure the FLAG constructs expressed protein of the correct size, infected and induced monolayers were lysed in reducing lane marker sample buffer and protein lysates were separated on 10% SDS-PAGE gels and transferred to a nitrocellulose membrane for western analysis of the FLAG-tagged protein. The membrane was blocked with PBS + 0.1% Tween 20 (PBS-T) and 5% nonfat milk prior to incubating in monoclonal anti-FLAG M2 antibody (1:40,000, Sigma, Thermo Scientific™) overnight at 4 °C followed by goat-anti mouse IgG-HRP secondary antibody (Invitrogen™) at room temperature for 2 hours. The membrane was developed with the Supersignal West Dura luminol and peroxide solution (Thermo Scientific™) and imaged using an Amersham Imager 600.

**Fig. S2.**
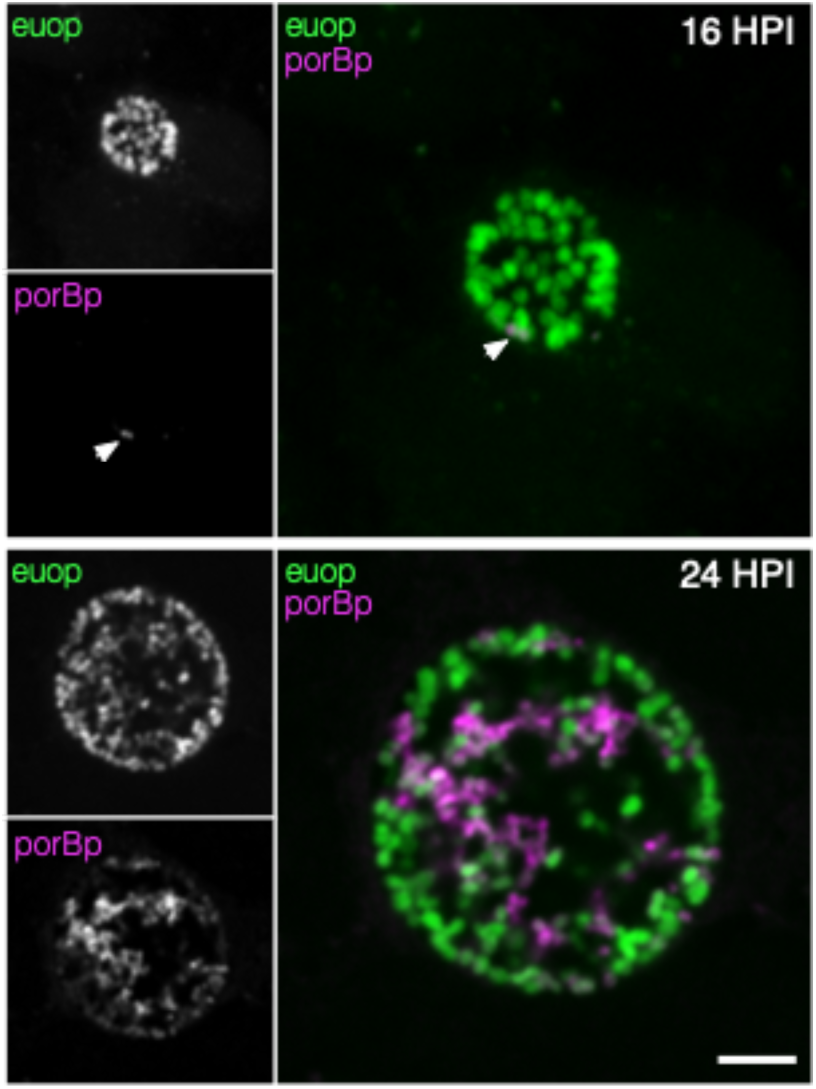
Cell type specific activity of the *porB* promoter. Cos-7 cells infected with the strain L2-PsciEng expressing Neongreen from the *euo* promoter (green) and Scarlet-I from the *porB* promoter (magenta). At 16 hpi there was only a single *porB* positive cell detected (arrow) while the rest of the chlamydial cells were only euoprom+. At 24 hpi there were two distinct cell populations, *euo*prom+ (green) and *porB*prom+ (magenta) cells. Size bar = 5µm.

**Fig. S3.**
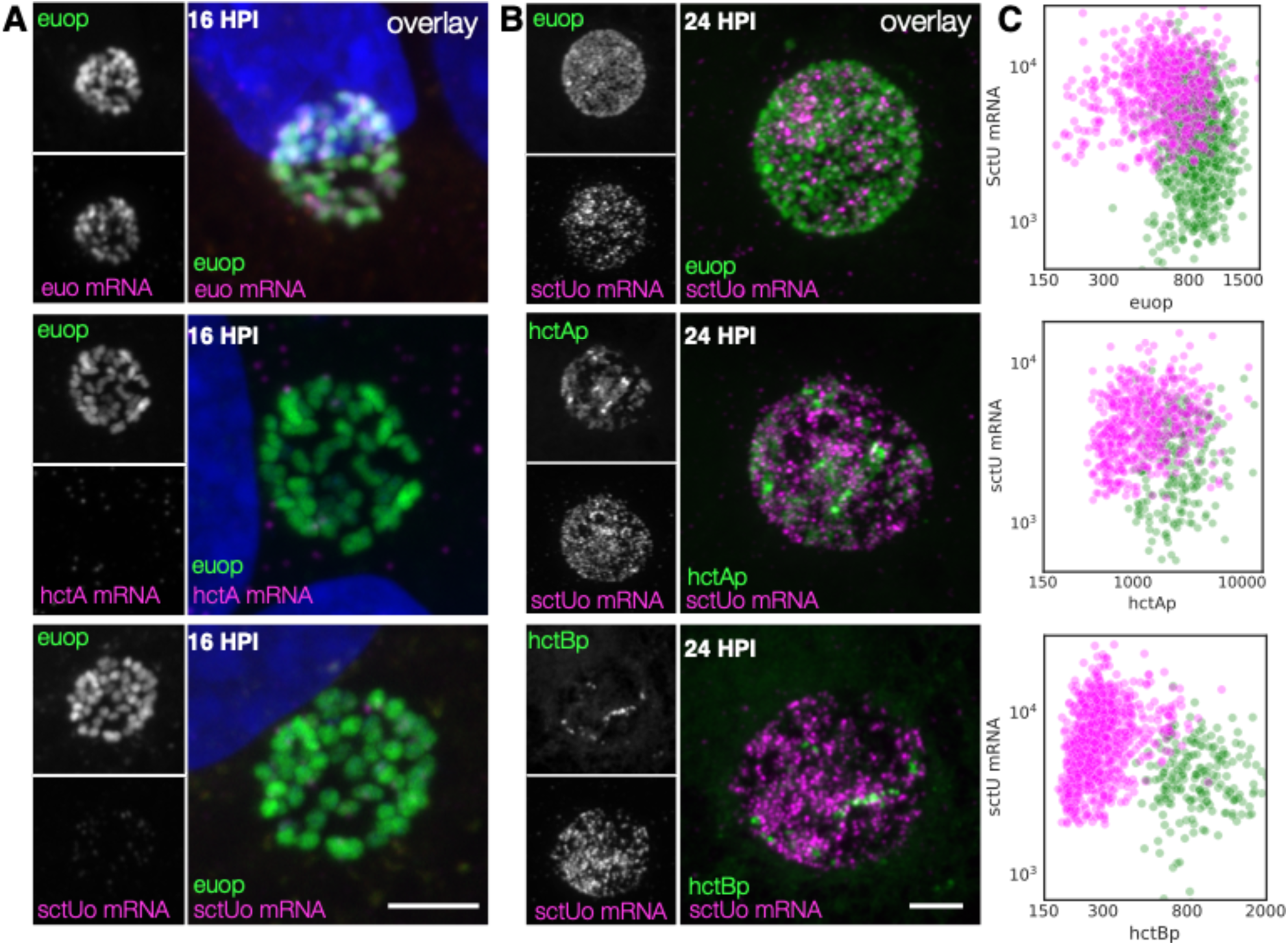
IB cell type expression of the T3SS structural operon *sctU*-op. (A) Cells were infected with L2-AsciEng for 16 hpi and fixed and stained using a FISH probe (*sctU* through *lcrD)* to the mRNA for the T3SS structural operon sctU-op. All cells were positive for *euo*prom expression (green) and negative for *sctU-op* mRNA (magenta). Infected cells were also probed for *hctA* mRNA expression and *euo* mRNA. Like *sctU*o the cells had little signal for the *hctA* mRNA. However, the *euo*prom+ cells were also positive for the *euo* mRNA (B) Cells were infected with L2-AsciEng and L2-BsciEng for 24 hpi and fixed and stained using FISH for the *sctU*-op mRNA. For the *euo*prom sample, the *sctU*-op FISH signal (magenta) was present in a distinct subset of cells and not in the majority of the *euo*prom+ cells (green). TrackMate was used to identify the *sctU*-op mRNA+ cells and the signal for *euo*prom and FISH were quantified for each *sctU*-op+ cell. The converse was also performed, the *euo*prom+ cells were identified (green) and the *euo*prom signal and FISH signal was quantified for each *euo*prom+ cell. The fluorescence intensity for each channel for both cell populations was plotted. The FISH signal was also compared to the *hctA*prom expression pattern and showed subsets of cells that were stained for both *sctU*-op mRNA and *hctA*prom expression as well as non overlapping populations. The *sctU*-op mRNA+ cells were again identified using TrackMate (magenta) and the signal for *hctA*prom and FISH were quantified for each *sctU*-op+ cell. Each *hctA*prom+ cell was also identified (green) and the FISH and *hctA*prom signal was determined and plotted for both cell populations. The *sctU-*op FISH staining was also compared to the expression from the *hctB*prom reporter. The *sctU*-op mRNA FISH staining was again present in a subset of cells but showed little overlap with the *hctB*prom fluorescent signal. The FISH signal and *hctB*prom signal were measured in both cell populations (*sctU* mRNA+ cells (magenta) and *hctB*prom+ cells (green) and plotted. Both populations were primary single positive, either *sctU*-op mRNA high or *hctB*rpom high but rarely both. Size bar = 5µm.

**Fig. S4.**
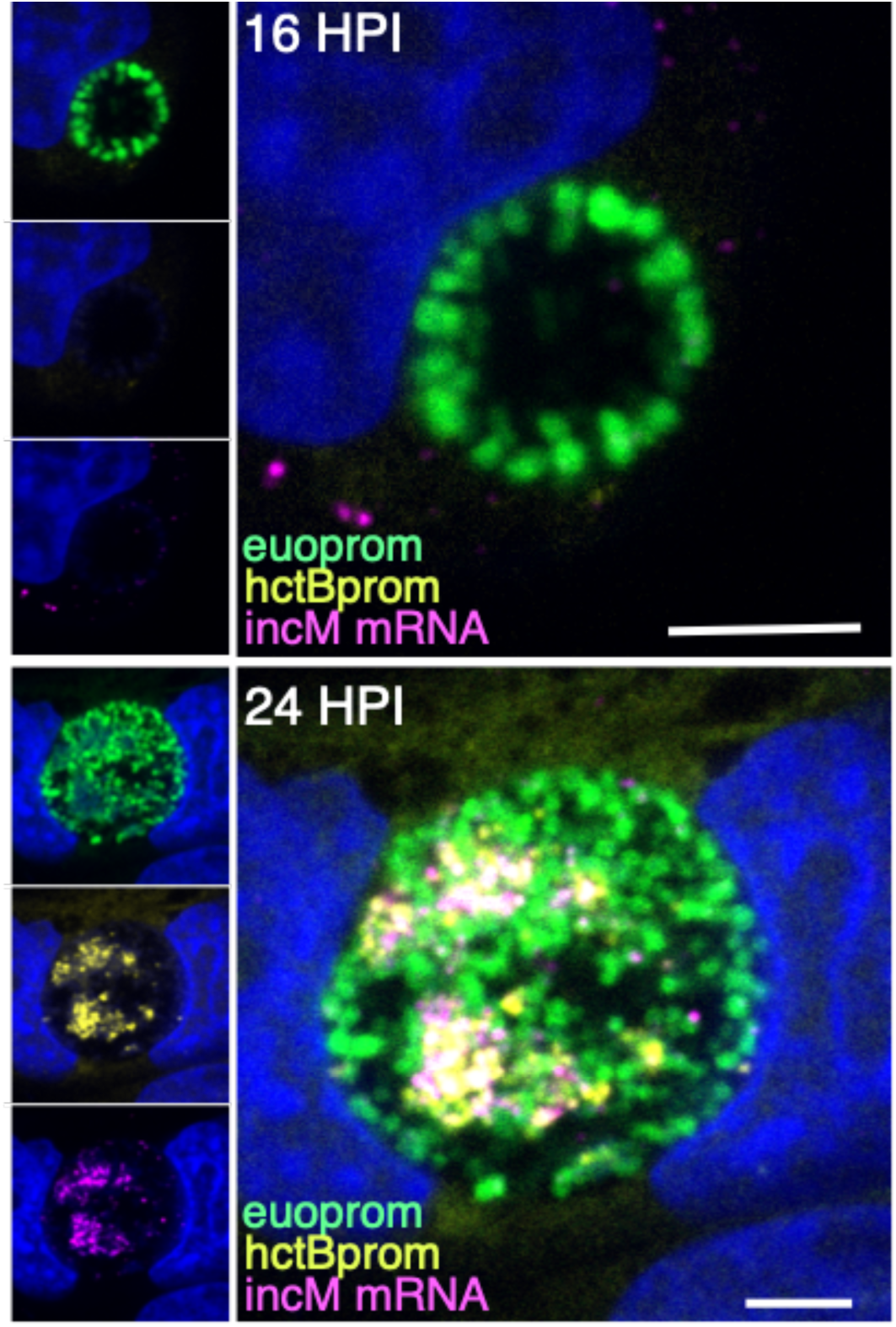
Cell type expression of *incM*. Cos-7 cells infected with L2-BsciEng for 16 and 24 hpi and stained for *incM* mRNA expression using custom FISH probes. The *incM* mRNA signal (magenta) was undetected at 16 hpi. At 24 hpi the *incM* mRNA signal showed overlap with the *hctB*prom signal (yellow) but not the *euo*prom signal (green). Size bar = 5µm.

