## Supplemental Table 2 for "The T3SS structural and effector genes of *Chlamydia trachomatis* are expressed in distinct phenotypic cell forms"

Quantification of single chlamydial expression plots

| Euo mRNA | Double postive | Single positive |
| --- | --- | --- |
| euo mRNA/hctAp | 7% | 93% |
| hctAp/euo mRNA | 7% | 93% |
| euo mRNA/hctBp | 1% | 99% |
| hctBp/euo mRNA | 5% | 95% |
| euo mRNA/euop | 90% | 10% |
| euop/euo mRNA | 91% | 9% |
| **HctB mRNA** | | |
| hctB mRNA/hctAp | 59% | 41% |
| hctAp/hctB mRNA | 52% | 48% |
| hctB mRNA/hctBp | 28% | 72% |
| hctBp/hctB mRNA | 61% | 39% |
| hctB mRNA/euop | 20% | 80% |
| euop/hctB mRNA | 18% | 82% |
| **HctA mRNA** | | |
| hctA mRNA/hctAp | 74% | 26% |
| hctAp/hctA mRNA | 77% | 23% |
| hctA mRNA/hctBp | 1% | 99% |
| hctBp/hctA mRNA | 32% | 68% |
| hctA mRNA/euop | 50% | 50% |
| euop/hctA mRNA | 28% | 72% |
| **PorB mRNA** | | |
| porB mRNA/hctAp | 56% | 44% |
| hctAp/porB mRNA | 78% | 22% |
| porB mRNA/hctBp | 3% | 97% |
| hctBp/porB mRNA | 20% | 80% |
| porB mRNA/euop | 70% | 30% |
| euop/porB mRNA | 73% | 27% |
| **SctJ mRNA** | | |
| sctJo mRNA/hctAp | 48% | 52% |
| hctAp/sctJo mRNA | 65% | 35% |
| sctJo mRNA/hctBp | 5% | 95% |
| hctBp/sctJo mRNA | 57% | 43% |
| sctJo mRNA/euop | 67% | 33% |
| euop/sctJo mRNA | 56% | 44% |

Quantification of single chlamydial expression plots cont.

| CTL0238o mRNA | Double postive | Single positive |
| --- | --- | --- |
| CTL0238o mRNA/hctBp | 3% | 97% |
| hctBp/CTL0238o mRNA | 11% | 89% |
| CTL0238o mRNA/euop | 88% | 12% |
| euop/CTL0238o mRNA | 71% | 29% |
| **Scc2o mRNA** | | |
| Scc2o mRNA/hctBp | 97% | 3% |
| hctBp/Scc2o mRNA | 89% | 11% |
| Scc2o mRNA/euop | 28% | 72% |
| euop/Scc2o mRNA | 10% | 90% |
| **incD mRNA** | | |
| incD mRNA/hctBp | 2% | 98% |
| hctBp/incD mRNA | 25% | 75% |
| incD mRNA/euop | 94% | 6% |
| euop/incD mRNA | 90% | 10% |
| **incV mRNA** | | |
| incV mRNA/hctBp | 98% | 2% |
| hctBp/incV mRNA | 60% | 40% |
| incV mRNA/euop | 24% | 76% |
| euop/incV mRNA | 3% | 97% |
